# Pentameric assembly of glycine receptor intracellular domains provides insights into gephyrin clustering

**DOI:** 10.1101/2022.11.10.512828

**Authors:** Arthur Macha, Nora Grünewald, Nastassia Havarushka, Nele Burdina, Christine Toelzer, Yvonne Merkler, Alfredo Cabrera-Orefice, Ulrich Brandt, Thomas Pauly, Luitgard Nagel-Steger, Karsten Niefind, Guenter Schwarz

**Affiliations:** Institute of Biochemistry, Department of Chemistry, University of Cologne, 50674 Cologne, Germany; The School of Biochemistry and Bristol Synthetic Biology Centre BrisSynBio, University of Bristol, Tankard’s Close, Bristol, BS8 1TD, UK; Cologne Excellence Cluster on Cellular Stress Responses in Aging-Associated Diseases (CECAD), University of Cologne, Germany; Radboud Institute for Molecular Life Sciences, Radboud University Medical Center, Nijmegen, The Netherlands; IBI-7, Structural Biochemistry, Forschungszentrum Jülich, 52425 Jülich & Institut für Physikalische Biologie, Heinrich-Heine-Universität Düsseldorf, 40225 Düsseldorf, Germany; Center for Molecular Medicine Cologne, University of Cologne, 50674 Cologne, Germany

## Abstract

Pentameric ligand-gated ion channels represent a large family of receptors comprising an extracellular domain, four transmembrane helices and a cytosolic intracellular domain (ICD). ICDs play important roles in receptor localization and trafficking, thus regulating synaptic activity and plasticity. Glycine and GABA type A receptor ICDs bind to the scaffolding protein gephyrin, a master regulator of inhibitory synapses. Here we report the use of yeast lumazine synthase as soluble pentameric protein scaffold for the study of receptor ICDs derived from GlyR α1− and β-subunits. We were able to create ICDs assemblies in a homo- (LS-βICD) and hetero-pentameric state (LS-αβICD) and provide first-in-class structural insights on their high structural flexibility using small angle X-ray scattering. We report a high-affinity interaction between the LS-αβICD and gephyrin leading to the *in vitro* formation of high-molecular mega-Dalton complexes composed of three gephyrin trimers and three pentamers as basic building block. Depending on the stoichiometric ratios between gephyrin and LS-ICDs the formed complexes grow or shrink in size. In cells, LS-ICDs efficiently recruited gephyrin and were able to accumulate gephyrin at GABAergic synapses in neurons. Our findings collectively propose a new, potentially general, mechanistic concept for a gephyrin-dependent bridging of GlyRs at the inhibitory synapse.

## Introduction

Pentameric ligand-gated ion channels (pLGICs) comprise excitatory and inhibitory receptors, which mediate efficient communication between neurons. All members of the pLGIC family, including the excitatory nicotinic acetylcholine receptor (nAchR), the inhibitory γ-aminobutyric acid receptor type A (GABA_A_R) and the glycine receptor (GlyR) share a similar subunit topology. An extracellular ligand-binding domain is followed by four transmembrane helices, of which the 3rd and 4th are connected by an enlarged intracellular cytoplasmic domain (ICD). Receptors of the pLGIC family can assemble either from five identical or different homologous subunits. The inhibitory GlyRs form homopentamers of α-subunits or heteropentamers composed of α- and β-subunits, with the α-subunit being present in one of three major splice forms (α1-α3). GABA_A_Rs, in contrast, are composed from a pool of up to 19 different subunits in mammals (Lemoine et al., 2012).

Once inserted in the plasma membrane, both inhibitory and excitatory neurotransmitter receptors diffuse laterally between synaptic and extrasynaptic compartments (Dumoulin et al., 2010, Chapdelaine et al., 2021). To ensure that at a given time a sufficient number of receptors is present at synapses, specific scaffolding proteins interact with the receptors and constrain their diffusion (Levi et al., 2008). Disruption of binding between receptors and corresponding scaffolding proteins has severe consequences on the balance between excitatory and inhibitory neurotransmission and may lead to hyperexcitability of the neuronal network. The latter often manifests in epileptic seizures, hyperekplexia and other types of neurological disorders (Mathews, 2007, Harvey et al., 2008, van Zundert et al., 2008, Olmos-Serrano et al., 2010, Dejanovic et al., 2015).

The synaptic confinement of inhibitory GlyRs as well as GABA_A_Rs mainly relies on the scaffolding protein gephyrin, which has been found in different oligomeric states. The assembly of gephyrin is an essential property for receptor clustering (Tyagarajan and Fritschy, 2014). Its oligomerization depends on the N-terminal trimeric G-domain and the C-terminal dimeric E-domain, which are evolutionary conserved and originate from the enzymatic function of gephyrin in basic metabolism (Schwarz et al., 2009, Sola et al., 2004). For its function at synapses, oligomers of gephyrin interact with receptors to build an extended postsynaptic receptor-scaffold network. It has been shown that the size of such a network and hence, the number of receptors located at the synapse, is controlled by the incorporation or dissociation of gephyrin molecules (Calamai et al., 2009).

The interaction between receptors and their scaffold is facilitated through the ICDs of pGLICs. It has been demonstrated that in case of GlyRs, only β-ICDs bind to gephyrin (Meyer et al 1995), whereas in GABA_A_Rs, a multitude of different subunit-ICDs mediate interactions with the scaffold (Tretter et al., 2008, Tretter et al., 2011, Kowalczyk et al., 2013). The core gephyrin-binding sequence within the GlyR β-ICD has been identified and encompasses 18 residues (Arg^394^-Ser^411^) (Meyer et al., 1995). Co-crystallization of the gephyrin E-domain (GephE) with the GlyR β-ICD peptide revealed an elongated binding site, which is primary dominated by hydrophobic interactions (Sola et al. 2004, Kim et al. 2006). Identification of the precise gephyrin-binding sequence fostered the application of isolated ICD-peptides for characterization of the interactions between inhibitory receptors and gephyrin. It has been demonstrated that GlyR β-ICD peptides bind to trimeric gephyrin via high- and low-affinity sites with dissociation constants (K_D_) in sub-micromolar and micromolar ranges, respectively (Herweg and Schwarz, 2012, Specht et al., 2011). GABA_A_R derived ICD-peptides were found to interact with gephyrin via the same binding site as GlyR β-ICD peptides, but with a approx. 100-fold lower affinity. This finding indicates a competition of the receptors for the binding site on gephyrin and a distinct binding mechanism between both receptor types and their common scaffold (Schrader et al., 2004, Specht et al., 2011, Tretter et al., 2008, Tretter et al., 2011, Kowalczyk et al., 2013, Maric et al., 2011, Maric et al., 2014).

Despite the broad application of the peptide-based approach for the characterization of their molecular interactions between receptor confining partners, this method carries certain disadvantages. Thus, it omits the physiological conformation of ICDs, neglects pentameric organization of the receptor and prohibits investigation of inter-subunits effects within heteromeric receptors (Herweg and Schwarz, 2012, Maric et al., 2011, Maric et al., 2014, Kowalczyk et al., 2013). To overcome those limitations, we established a novel model system, which enables the attachment of the full-length ICDs to a pentameric ‘platform’ thus mimicking the physiological oligomerization and folding of the scaffold-interacting domains of pLGICs. For this purpose, we utilized a soluble protein from *Saccharomyces cerevisiae* - lumazine synthase (LS), which exhibits spatial dimensions similar to the pentameric receptors of the pLGIC family. We incorporated α1-ICD and β-ICD of the GlyR subunits into LS, which assembled into stable complexes with a two to three stoichiometry. We demonstrated β-ICD-directed high-affinity interactions of LS-variants with gephyrin *in vitro* and in cultured neurons, promoting the formation of high-molecular weight complexes. This enabled us to utilize small angle scattering-based structural analysis to disclose several ICD conformations, indicating orchestrated conformational changes of GlyR-ICDs within a pentameric assembly. Finally, we simulate receptor-scaffold clustering *in vitro* by using a combination of various analytical methods, providing a new molecular mechanism underlying postsynaptic clustering. In summary, this study introduces LS as universal model system for the characterization of ICDs of all types of pLGIC receptors thus allowing the identification of the general principle of receptor clustering at postsynaptic sites.

## Results

### Biochemical characterization of LS-ICD chimeras

LS forms a spherical, flattened molecule with a diameter similar to those of pentameric pLGIC receptors (Figure 1, Table S1). Each LS monomer consists predominantly of α-helices and harbors a short surface-exposed α4-α5 loop (Figure 1a and b), which shows a distance to the loops of neighboring subunits of 27 Å on average (Figure 1b). This loop-to-loop distance compares well to the distance of 25.7 Å, observed between the terminal ends of ICDs in the structure of the GlyRα_1_ receptor (Figure 1c) (Du et al., 2015). Therefore, the entire ICDs of the GlyR α1-subunit (residues 338-427) and β-subunit (residues 334-454) were introduced into LS, forming chimeric constructs LS-αICD and LS-βICD, respectively. Both LS variants were expressed as His-tagged proteins in *E. coli*, either separately to produce homopentameric LS-αICD or LS-βICD proteins or co-expressed to generate LS-αβICD hetero-oligomers. Unmodified LS (LS^wt^) was used as a control throughout this study.

**Figure 1.**
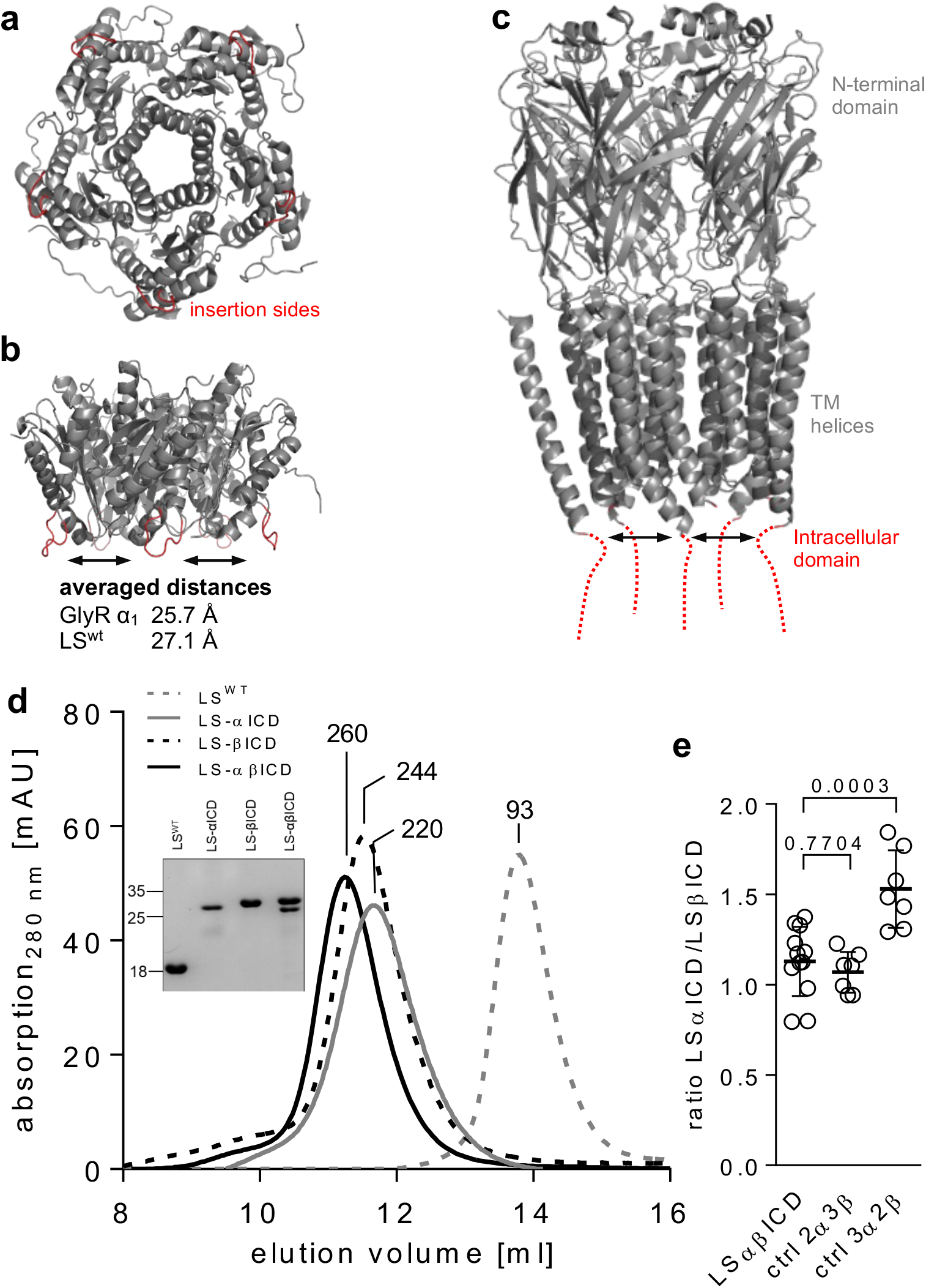
Structural similarity of pentameric LS and GlyR α1. **(a)** Top view of crystal structure of pentameric yeast lumazine synthase (PDB: 1EJB) with highlighted insertion sites for GlyR-ICD between α-helices 4 and 5 are highlighted in red. **(b)** Side view of lumazine synthase showing the insertion sites pointing downwards. **(c)** Cryo-EM structure of GlyR α1 (PDB: 3JAF). N-terminal domain and transmembrane helices are shown in grey, with indicated position of cytoplasmic helices of ICDs (red). Averaged distances between insertion sites or cytoplasmic loops within a pentameric LS or GlyR were determined using Pymol. **(d)** Representative elution profiles of LS-ICD chimeras and LS^wt^. Determined MWs [kDa] of LS chimeras according to standard protein calibration curve are indicated. Insert shows SDS-PAGE analysis of respective SEC fraction of LS variants obtained after SEC. **(e)** Densitometric determination of LS-βICD and LS-αICD-ratio in purified LS-αβICD. Data were analyzed for statistical significance using a one-way ANOVA with multiple comparison test. Exact p values are indicated.

By applying affinity purification and size exclusion chromatography (SEC), LS^wt^ and LS GlyR-ICD variants were enriched to high homogeneity, as judged by SDS-PAGE analysis (Figure 1d). LS^wt^, LS-αICD and LS-βICD showed distinct bands with apparent molecular masses close to the expected values of 19.9 kDa, 30.0 kDa and 32.9 kDa, respectively, thus confirming successful incorporation of the GlyR-ICDs into the LS backbone. Purified LS-αβICD yielded two distinguishable bands, demonstrating the integration of both LS-αICD and LS-βICD upon co-expression into one stable assembly. SEC demonstrated a molecular weight (MW) of 93 kDa for LS^wt^, which is in a close agreement to the predicted mass of 99 kDa for a LS^wt^ pentamer. In contrast, the ICD-carrying LS-variants eluted with molecular masses exceeding the expected size of pentamers of 150-165 kDa. LS-αICD and LS-βICD elution corresponded to molecular masses of 220 and 244 kDa, respectively, suggesting an increased hydrodynamic radius due to a more extended conformation of LS-ICD chimeras in comparison to the globular LS^wt^. LS-αβICD showed the lowest elution volume corresponding to 260 kDa, indicating different inter-subunit contacts as compared to the respective homo-pentameric assembly.

To determine the MWs of LS variants more accurately we applied analytical ultracentrifugation (AUC). By utilizing velocity equilibrium experiments, MWs were determined for LS^wt^, LS-βICD and LS-αβICD, which matched the expected MW of the respective LS variants in a pentameric assembly (Figure S1a-e). However, for LS-αICD an increased MW of 258 kDa was determined, which was caused by self-aggregation as confirmed with its progressive sedimentation at higher centrifugal velocities (Figure S1b and e).

To further demonstrate that the structural integrity of LS backbone has been maintained in all LS variants, we determined their riboflavin content by UV/vis spectroscopy showing similar absorbtion at 410 nm (Figure S1f). In addition, CD spectroscopy disclosed ICD-specific differences between LS^wt^ and the LS-ICD variants suggesting a specific folding of the corresponding ICDs (Figure S1g). Taken together, our biochemical analyses collectively confirmed that introduction of the GlyR-ICD into the LS pentamer resulted in the formation of pentameric complexes that were stable for further biochemical analyses.

In light of the different reports on subunit composition in heterotetrameric GlyRs we aimed to determine the stoichiometry of the ICD-subunit within LS-αβICD by densitometric quantification of Coomassie blue-stained bands following SDS-PAGE. By utilizing homomeric LS-αICD and LS-βICD as standard proteins, the obtained band intensities of LS-αβICD were translated to total protein amount (Figure S1h-j). To further assess the determined ratio, two control samples with distinct ratios of either 2 LS-αICD / 3 LS-βICD or 3 LS-αICD / 2 LS-βICD were utilized for comparison. As a result, the 2 LS-αICD/ 3 LS-βICD control showed a similar ratio as the LS-αβICD pentamer, demonstrating a 2αICD/3βICD stoichiometry within LS-αβICD (Figure 1e). In addition, thermostability of LS-ICD chimeras using differential scanning fluorimetry (DSF) and CD-spectroscopy (Figure S1k-n) yielded LS-αβICD as most stable complex with a T_m_ of 51 °C, which indicates additional stabilizing interactions (in comparison to the homopentamers) between αICD and βICD within the heteropentamer.

### Small angle X-ray scattering provides structural insights on GlyR ICDs

Up to date, structural information of the ICDs of any member of the pLGICs receptor family remains unknown (Huang et al., 2015, Du et al., 2015, Unwin, 2005, Miller and Aricescu, 2014, Kumar et al., 2020). To decipher the structure of GlyR-ICDs, LS-βICD and LS-αβICD were subjected to small-angle X-ray scattering (SAXS), which allows to access low-resolution structures in solution.

Based on the scattering data of LS-βICD and LS-αβICD, the molecular masses of 163 kDa and 174 kDa were determined, respectively, confirming the expected molecular masses of 164 kDa (LS-βICD) and 159 kDa (LS-αβICD), respectively, within acceptable limits of error (Table 1). The scattering of LS-βICD and LS-αβICD exhibited a linear regression in the Guinier region, supporting a high homogeneity of the analyzed samples and the absence of protein aggregates (Figure 2a). Differently from LS^wt^, the profile of the *P*(*r*) function for LS-βICD and LS-αβICD was asymmetrical with an extended shoulder, suggesting that both LS-βICD and LS-αβICD formed rather elongated than spherical particles (Figure 2b). This was consistent with the significantly higher *D*_max_ obtained for LS-βICD and LS-αβICD (Table 1), in comparison to the theoretical values of 87.1 Å and 85.9 Å, respectively, which were expected for spherical proteins with the same number of residues as LS-βICD (1,500) and LS-αβICD (1,438) (Durand et al., 2010).

**Table 1.** SAXS-derived parameters of LS^wt^ and LS-ICD variants.

|  | LS<sup>wt</sup> | LS-βICD | LS-αβICD |
| --- | --- | --- | --- |
| **Structural parameters** |  |  |  |
| $I(0)$ from $P(r)$ | $61.38 \pm 0.02$ | $86.53 \pm 0.07$ | $79.20 \pm 0.04$ |
| $R_g$ from $P(r)$ (nm) | $2.96 \pm 0.01$ | $4.69 \pm 0.01$ | $4.44 \pm 0.01$ |
| $s$-range for Guinier fit ($\text{nm}^{-1}$) | 0.19 - 0.42 | 0.15 - 0.27 | 0.07- 0.23 |
| $I(0)$ from Guinier fit | $61.57 \pm 0.04$ | $86.62 \pm 0.08$ | $79.45 \pm 0.05$ |
| $R_g$ from Guinier fit, Auto$R_g$ (nm) | $2.99 \pm 0.03$ | $4.65 \pm 0.13$ | $4.44 \pm 0.04$ |
| $D_{\text{max}}$ (nm) | 9.00 | 16.72 | 15.50 |
| POROD volume estimate ($\text{nm}^3$) | $149 \pm 15$ | $355 \pm 20$ | $334 \pm 15$ |
| **Molecular mass (kDa)** |  |  |  |
| From sequence | 99.5 | 164.2 | 158.5 |
| From SAXS data | 106 | 163 | 174 |

**Figure 2.**
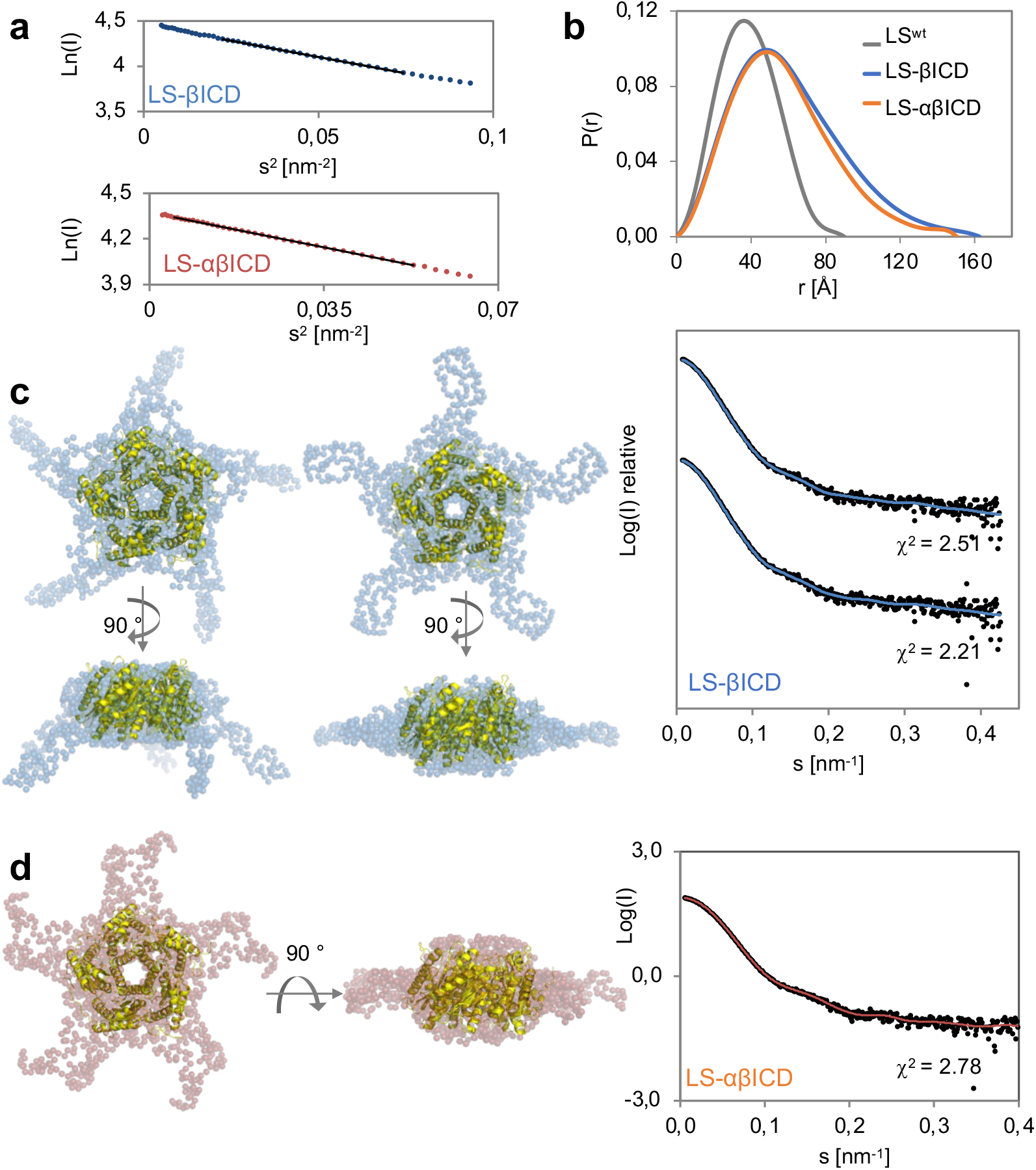
SAXS analysis and *ab initio* GASBOR models of LS-βICD and LS-αβICD. **(a)** Linear fit (black lines) of the LS-βICD and LS-αβICD scattering curves (blue and red dots, respectively) within Guinier region. **(b)** P(r) function of the LS-βICD and LS-αβICD scattering curves in comparison to LS^wt^. **(c-d)** Representative P5 GASBOR models of LS-βICD (blue) and LS-αβICD (red) superimposed with LS crystal structure (yellow). Water molecules were omitted for clarity. On the right – corresponding fits of the LS-βICD and LS-αβICD GASBOR models (blue and red lines) to the experimental intensities (black dots). Goodness-of-fit (χ2) values obtained for each model are shown.

*Ab initio* modelling with GASBOR imposing P5 symmetry constrains resulted in one type of conformation for both of LS-βICD and LS-αβICD, showing a clearly defined LS-core and GlyR-ICDs emerging from its sides as loop-like structures and positioned at the same plane as the LS-backbone or slightly bent down (Figure 2c and d). Models generated with a second program DAMMIF (P5 symmetry) confirmed the previous data and disclosed additional conformations with ICDs positioned together (Figure S2a and b). Our results are consistent with the predicted flexibility of GlyR-ICD. Consequently, LS-αβICD and LS-βICD could occur in several asymmetrical and symmetrical intermediate states. Thus, the observed final scattering curve of both LS variants would represent an average of scattering profiles of different conformers. Solely upon application of a 5-fold symmetry constrain, two extreme states of LS-αβICD and LS-βICD become accessible: the one, where ICDs are entirely detached from each other -forming an “open” conformation, and the other, where all five ICDs are in close proximity – representing a “closed” state (Figure 2c and d and Figure S2a and b).

The flexibility of GlyR-ICDs within LS-βICD and LS-αβICD was further supported by a dimensionless Kratky plot of the scattering data, with LS-βICD and LS-αβICD maxima shifted to higher values while the function became more extended with increasing R_g_×s (Figure S2c). Thus, the hypothesis was supported that a considerable degree of flexibility has been introduced into LS-βICD and LS-αβICD due to the presence of the ICDs (Receveur-Brechot and Durand, 2012).

### Gephyrin binds to LS-ICD chimeras with high affinity

The formation of heteromeric GlyR receptors is a prerequisite for the gephyrin-dependent localization of GlyRs at the synapse. Therefore, we examined binding of LS-ICDs chimeras to gephyrin using different methods (Figure 3 and 4). First, the interactions of LS-βICD and LS-αβICD with purified GephE dimer as well as trimeric gephyrin were analyzed by isothermal titration calorimetry (ITC) (Figure 3a and b). As control, gephyrin or GephE were injected into LS-αICD and as expected, no binding was observed (Figure S3a and b), verifying the absence of any unspecific interaction between the LS backbone as well as αICD and gephyrin.

**Figure 3.**
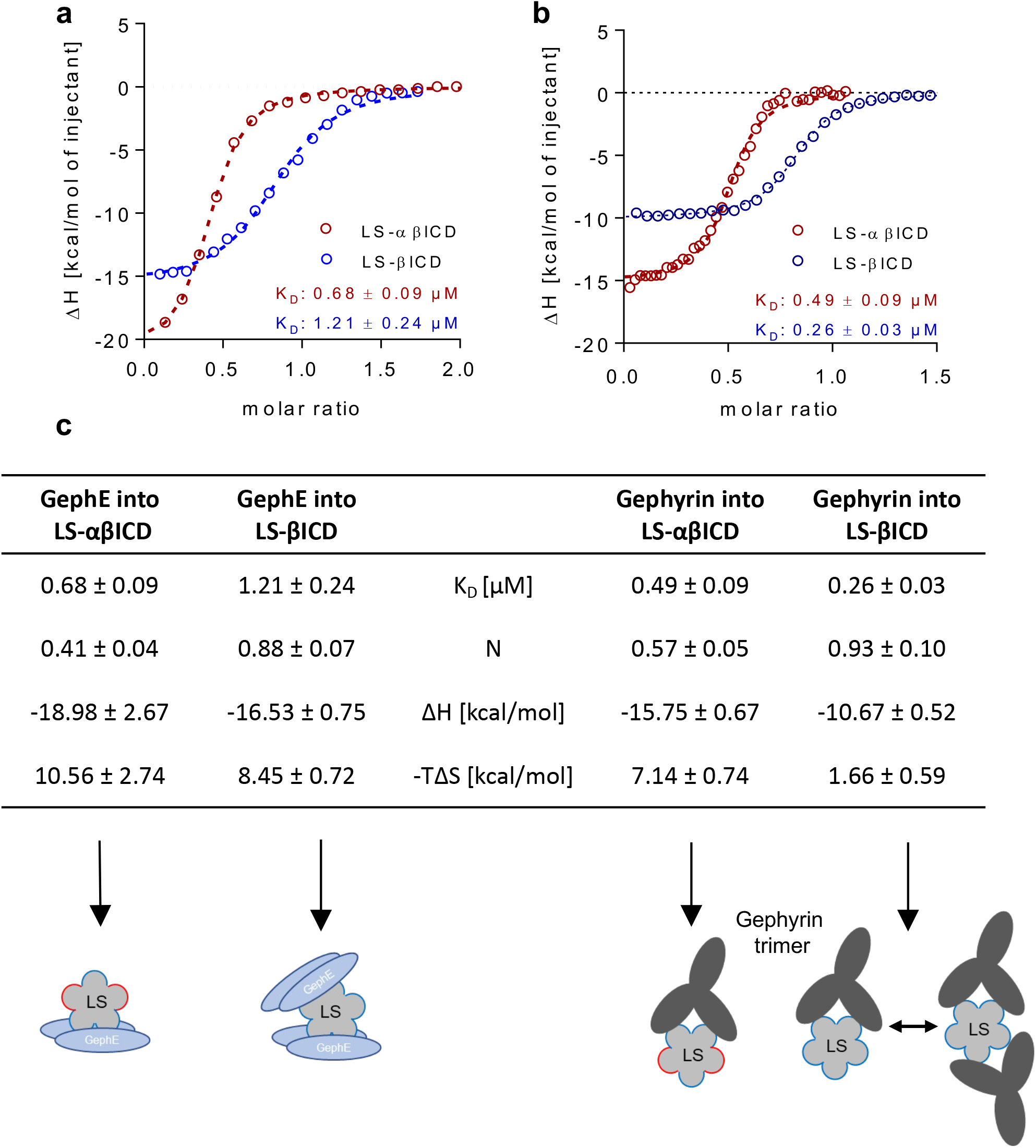
Interaction of LS-ICD variants with gephyrin. **(a-b)** Representative binding isotherms (dots) with one site binding fit (dotted line) of LS-αβICD (red) or LS-βICD (blue) titrated with GephE (a) or titrated with gephyrin (b). **(c)** Summary of binding parameters obtained from respective ITC experiments showing values for dissociation constant K_D_ [μM], stoichiometry N, binding enthalpy ΔH [kcal/mol] and binding entropy –TΔS [kcal/mol]. Depicted values presented as mean ± SD, n = 3. (c, lower panel) Descriptive models visualizing complex formations of LS with GephE or with gephyrin trimer based on the stoichiometries derived from ITC experiments.

**Figure 4.**
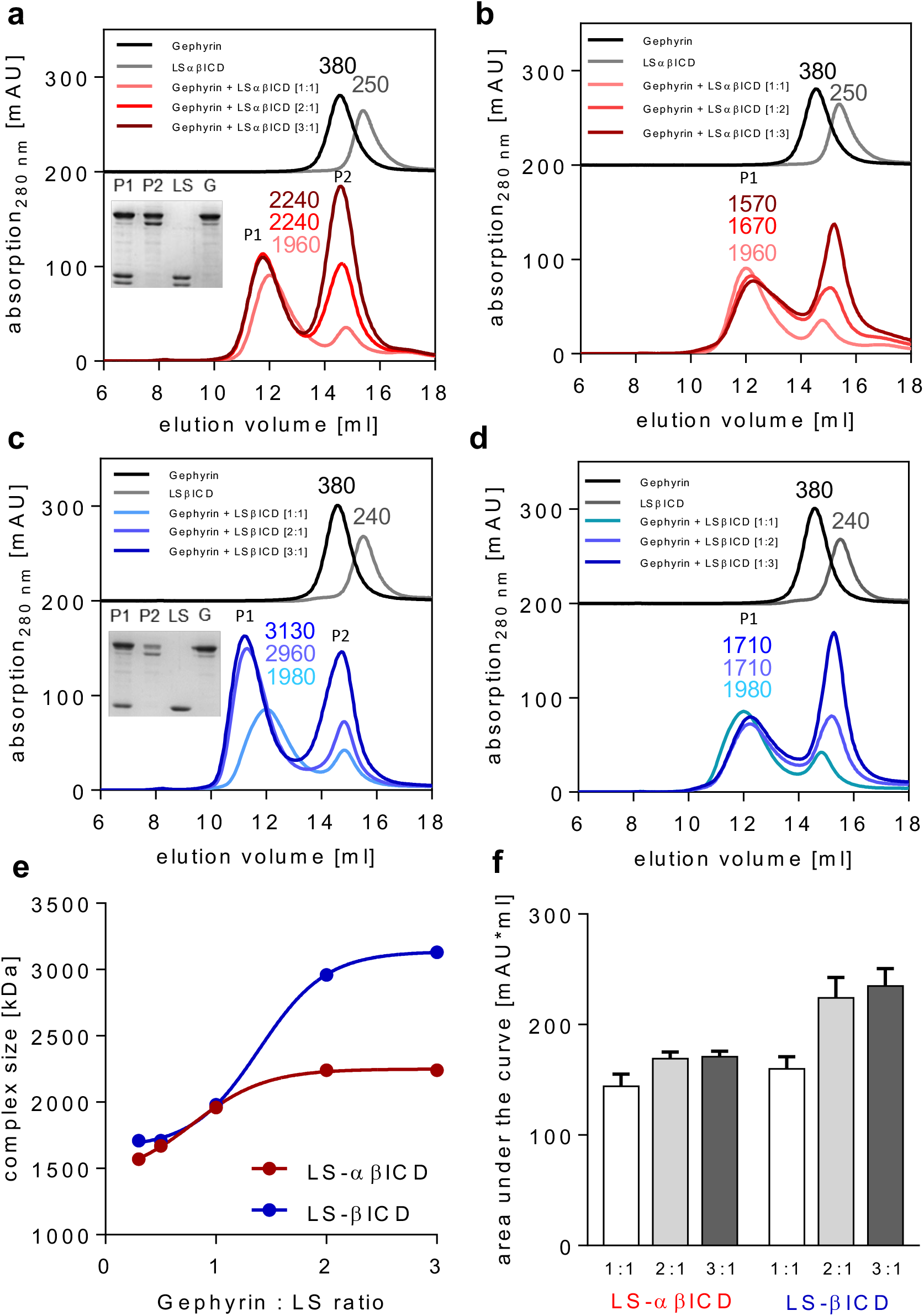
Molecular weight determination of LS-ICD-gephyrin complexes. **(a-d)** Representative elution profiles of gephyrin with LS-αβICD (red) and gephyrin with LS-βICD (blue) at equimolar protein concentrations (1:1) or with two-fold (2:1) or three-fold (3:1) gephyrin excess (a, c) or with two-fold (1:2) and three-fold (1:3) LS excess (b, d). Inserts in (a) and (c) depicting SDS-PAGE analysis of peak 1 (P1) and peak 2 (P2) with gephyrin (G) and LS-αβICD or LS-βICD (LS) as reference, respectively. Numbers indicate determined MW [kDa] of proteins. Single gephyrin trimer (black) and LS-ICD pentamers (grey) served as reverence. **(e)** Determined complex MWs (P1) from a-d plotted against protein ratios of the respective SEC experiment with applied sigmoidal fit to visualize graph propagation. **(f)** Determined areas under the curve of peak 1 (P1) from (a) and (c). Data presented as mean ± SD, n = 3.

Next, ITC experiments were performed by injecting purified GephE to LS-αβICD or LS-βICD. Isotherms were fitted to a one-site binding model to obtain binding parameters (Figure 3a and c and S3c and d). Both experiments resulted in an exothermic binding event with dissociation constants of K_D_ = 0.68 ± 0.09 μM for LS-αβICD and K_D_ = 1.12 ± 0.24 μM for LS-βICD and molar ratios of N = 0.41 ± 0.04 and N = 0.88 ± 0.07 for LS-αβICD and LS-βICD, respectively. The determined stoichiometries suggest a saturation of LS-αβICD with 1 GephE dimer and LS-βICD with 2 GephE dimers (Figure 3c, lower panel). Both interactions showed enthalpy-driven characteristics with comparable entropically contributions.

When injecting full-length gephyrin to either LS-αβICD or LS-βICD, again exothermic signals were detected (Figure 3b and c, S3e and f). Obtained isotherms were fitted using a one-site binding model and dissociation constants of *K*_D_= 0.49 ± 0.09 μM and *K*_D_ = 0.26 ± 0.03 μM were derived for LS-αβICD and LS-βICD, respectively (Figure 3c). This finding suggests that in both ICD chimeras, the presence of LS-βICD mediates high-affinity interaction with gephyrin, irrespective of the presence of the αICD in LS-αβICD. For both LS variants a strong negative enthalpic term (ΔH) together with a positive entropic term (-TΔS) was observed, indicating an enthalpy-driven binding between LS-βICD or LS-αβICD and gephyrin (Figure 3c).

The stoichiometry derived from the binding isotherms was analyzed in respect to the number of possible interaction sites in LS-αβICD and LS-βICD pentamer (Figure 3c). In that sense, the determined stoichiometry of N = 0.57 ± 0.05 for LS-αβICD and N =0.93 ± 0.1 for LS-βICD translate to a saturation of LS-αβICD with 1 gephyrin trimer and LS-βICD with 1–2 gephyrin trimers (Figure 3c, lower panel). To further understand the nature of this interaction, we next determined the molecular weights of the gephyrin–LS-αβICD/ LS-βICD complexes.

### Gephyrin binding to LS-αβICD and LS-βICD yielded distinct high-molecular complexes

*In vitro* complex formation of LS variants with gephyrin was further analyzed via analytical SEC (Figure 4). To ensure a high saturation of binding sites, we used 40-120 μM of either gephyrin or LS-variants, which was an approx. two orders of magnitude higher concentration than the ITC-determined K_D_ values of their respective interaction. As a matter of control, LS-αICD and LS^wt^ were applied separately as controls and showed no binding to gephyrin (Figure S4a and b).

In the first titrations, LS chimeras were kept constant at 40 μM and co-incubated with 40-120 μM gephyrin. Both LS-αβICD and LS-βICD formed complexes with gephyrin as seen by a shift of the elution volumes towards high-molecular weights in comparison to the individual proteins and the sum of both individual proteins (Figure 4a and c). Following equimolar interaction, a single additional peak with a MW of 1,960 kDa for LS-αβICD was observed (Figure 4a, P1). The absence of any unbound LS-αβICD and the low amount of remaining gephyrin (Figure 4a, P2) suggests a complete binding of the entirety of LS pentamers to gephyrin. When increasing the amount of gephyrin in relation to LS-αβICD by two (2:1) and three-fold (3:1) molar excess in comparison to the respective monomers, the MW of the resulting complexes increased to 2,240 kDa. As both conditions produced a similar complex in size, we assumed that a two-fold excess of gephyrin was sufficient to maximize the size of the gephyrin-LS-αβICD complex. In reverse, when raising the LS-αβICD concentration by two or three-fold, respectively, while leaving the gephyrin concentration constant at 40 μM, no increase in MWs of the resulting complexes were detected (Figure 4b). The MW rather decreased to 1,570 kDa with 3-fold excess of LS-αβICD and exhibited a tailing shoulder, suggesting an increase polydispersity of the complex (Figure 4b). The different complexes formed are summarized in Figure 5.

**Figure 5:**
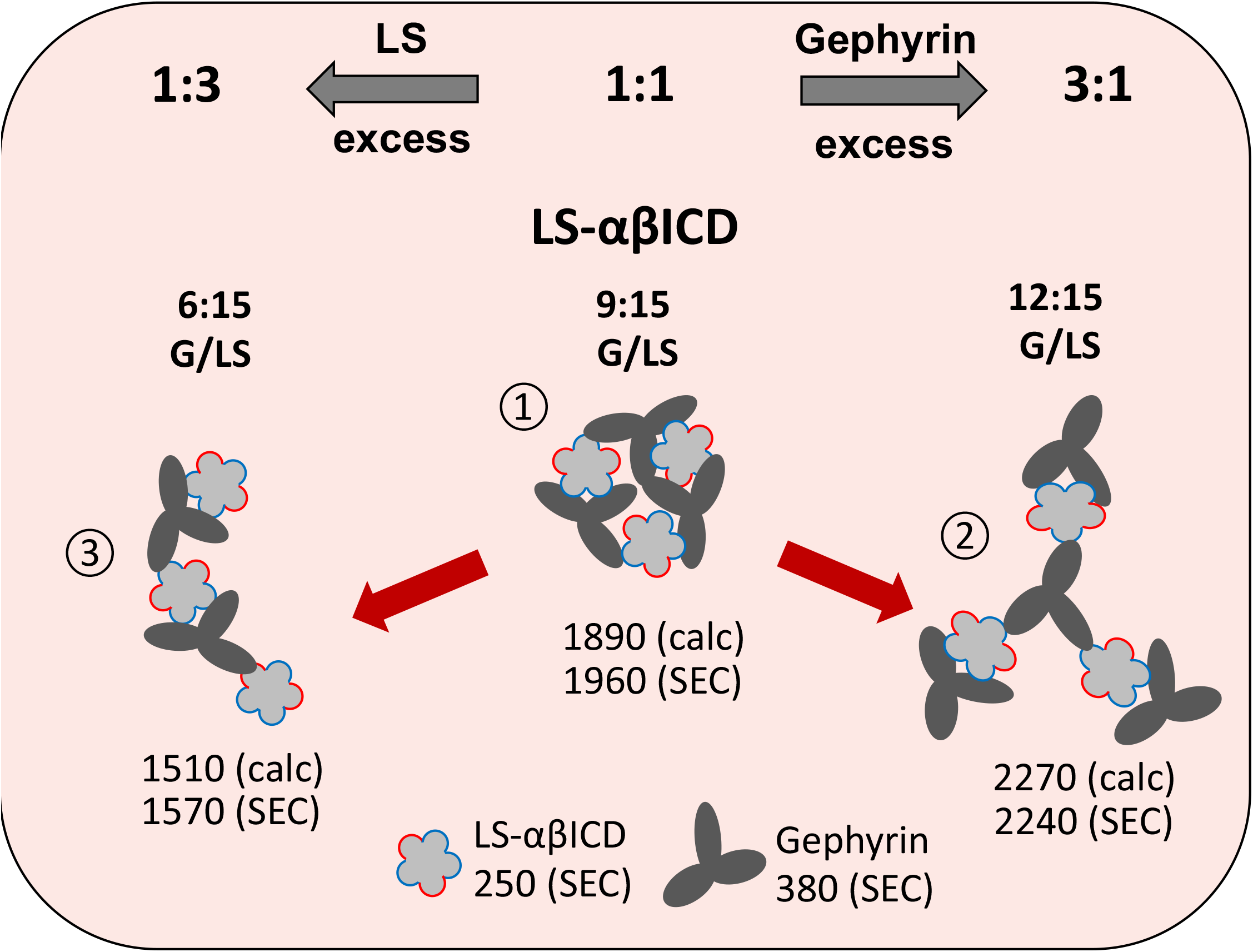
Model of gephyrin-LS-αβICD complex formation. Depicted are the respective complexes formed at equimolar quantities (1:1) and with three-fold gephyrin (3:1) or LS (1:3) excess. In a 1:1 equimolar setup LS-αβICD variant was saturated with the same amount of gephyrin molecules, resulting in a complex of three gephyrin trimer and three LS pentamers with MWs of 1960 kDa (1) Utilizing gephyrin excess led to an enlargement of the complex by one gephyrin trimer, increasing the MW to 2240 kDa (2). LS excess resulted in the formation of smaller complexes through the release of gephyrin molecules, probably due to the oversaturation of gephyrin binding sites (3).

Co-incubation of gephyrin with LS-βICD in an equimolar manner resulted again (similar to LS-αβICD) in a complete binding of LS-βICD in a high molecular mass complex of 1,980 kDa with only trace amounts of unbound gephyrin left (Figure 4c, P1 and P2). Following an increase of gephyrin by two- and three-fold molar excess to LS-βICD, a strong and significant gain in MW saturating at 2,960 kDa (2:1) and 3,130 kDa (3:1) was observed (Figure 4c). In analogy to LS-αβICD, the excess of LS-βICD relative to gephyrin did not result in the formation of larger protein complexes and the resulting gephyrin-LS-βICD species decreased in size to 1,710 kDa at a 1:2 and 1:3 ratio respectively (Figure 4d and Figure S5).

While both gephyrin-LS chimera complexes revealed a similar MW in the presence of LS excess (1,570-1,710 kDa), complexes with different sizes were formed when gephyrin was provided in excess (Figure 4e). Notably, the MW of gephyrin-LS-βICD reached a size (3,130 kDa) being almost 50% larger than gephyrin-LS-αβICD (2,240 kDa), suggesting the binding of more gephyrin molecules to LS-βICD (Figure 4e). When comparing the different peak areas in both titrations (LS-αβICD and LS-βICD, Figure 4f), a clear gain in peak size was observed for the gephyrin-LS-βICD complexes being compatible with more gephyrin molecules bound to the complex. In reverse, when inspecting the amounts of remaining unbound gephyrin, we found 3–5.25 nmol gephyrin bound to 4 nmol LS-βICD with increasing gephyrin quantity (Figure S4c and d). In contrast, areas of LS-αβICD complexes showed little gain with gephyrin excess and the amount of bound gephyrin increased moderately from 3 nmol to 4 nmol gephyrin (Figure 4f and Figure S4d).

In aggregate, we conclude that both LS-βICD and LS-αβICD bound 75% of gephyrin under conditions of equimolar interaction, which would translate to the binding of one gephyrin trimer to one LS pentamer. Given the determined MW of the individual proteins in SEC (Figure 4a-d), the detected masses fit well to an interaction of three gephyrin trimers with three LS-αβICD / LS-βICD pentamers. Note, that multiples of individual masses as determined by SEC are 1,890 kDa and 1,860 kDa, respectively. With gephyrin excess, more gephyrin molecules were recruited, resulting in a complex consisting of four gephyrin trimers and three LS-αβICD pentamers (2,270 kDa) as well as six gephyrin trimers and three LS-βICD pentamers (3,000 kDa), respectively. In both cases, the detected complexes require multiple interactions between gephyrin and the ICDs of different LS molecules (Figure 5 and S5).

### LS-ICD chimeras are recruited to gephyrin clusters

Based on our *in vitro* binding and complex formation studies, we next aimed to proof that LS-ICDs are able to form similar type of interaction in non-neuronal and neuronal cells. First, mCherry-gephyrin was expressed in COS-7 cells (Figure 6a-d) resulting in the formation of well-known aggregates, so-called gephyrin *blobs* (Kirsch et al., 1995, Saiyed et al., 2007). Upon co-transfection of mCherry-gephyrin with GFP-tagged LS^wt^ and LS-αICD (lacking β-ICD), both LS variants showed either diffusive cytoplasmic distribution or nuclear staining and no co-localization with gephyrin *blobs* (Figure 6a and b). In contrast, co-expression of mCherry-gephyrin and LS-βICD-GFP resulted in retrieval of LS-βICD-GFP into gephyrin *blobs* (Figure 6c), indicating a specific interaction of the β-ICD and gephyrin (Figure S6). In the same manner, triple transfection of COS-7 cells with LS-αICD-GFP, LS-βICD-HA and mCherry-gephyrin resulted in the relocation of the LS-αICD-GFP signal into gephyrin *blobs* (Figure 6d). This finding confirms that LS-αICD-GFP and LS-βICD-HA formed stable heteropentamers, which were able to undergo β-ICD-directed interaction with gephyrin in COS-7 cells.

**Figure 6.**
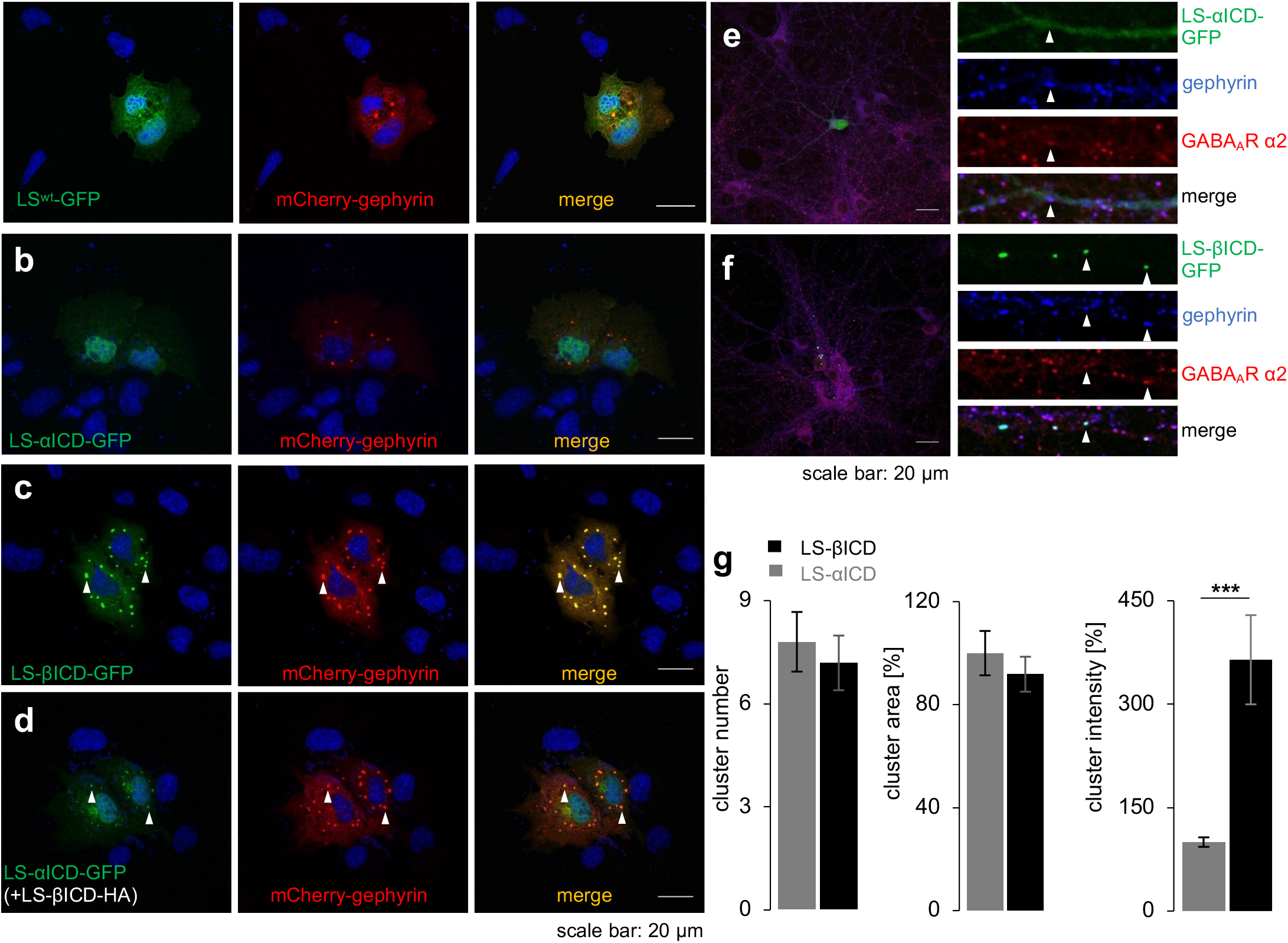
Interaction between gephyrin and LS variants in cells. **(a–d)** Co-expression of mCherry-gephyrin in COS7 cells with (a) LS^wt^-GFP, (b) LS-αICD-GFP, (c) LS-βICD-GFP, and (d) LS-αICD-GFP and LS-βICD-HA. Gephyrin blobs are highlighted with arrow heads. **(e, f)** PHNs were transiently transfected to express LS-αICD-GFP (e) or LS-βICD-GFP (f) and further immunostained for endogenous gephyrin (blue), together with GABAAR α2 (red) to visualize inhibitory synapses. A section is represented in higher magnification to highlight the gephyrin signal. Gephyrin/GABAAR α2 puncta are shown as arrow heads. **(g)** Quantification of gephyrin cluster number, area and intensity within neurons expressing LS-αICD-GFP or LS-βICD-GFP via two-tailed t-test from n=3 independent experiments.

Next, the interaction between LS variants and gephyrin was analyzed in dissociated primary hippocampal neurons DIV14 (Figure 6e-g). Inhibitory synapses were visualized through the overlap of *punctate* immunoreactivity of α_2_-subunit-containing GABA_A_Rs with that of gephyrin (Figure 6e-f). In hippocampal neurons, transfected LS-αICD-GFP was found to be diffusively distributed within the soma and dendrites of the neurons and showed no co-localization with endogenous gephyrin clusters (Figure 6e). In contrast, LS-βICD-GFP was accumulated at gephyrin-positive *puncta* at inhibitory synapses (Figure 6f). The influence of LS-βICD-GFP on gephyrin clustering was further quantified and compared to the non-interacting variant LS-αICD-GFP. Neither LS-βICD-GFP nor LS-αICD-GFP affected the number and area of gephyrin clusters (Figure 6g). However, in comparison to LS-αICD-GFP, expression of LS-βICD-GFP resulted in a highly significant (P < 0.001) four-fold increase of gephyrin cluster intensity (Figure 6g). This finding demonstrates the ability of LS-βICD to (i) bind to synaptic gephyrin clusters and (ii) to recruit additional gephyrin molecules to synaptic sites without changing the overall size of the GABAergic synapse. Thus, this result is in line with our observed *in vitro* formation of high-molecular complexes between gephyrin and LS-βICD that involved several gephyrin and LS-ICD oligomers.

## Discussion

The number of neurotransmitter receptors at the postsynaptic site defines the strength of a synapse. Lateral diffusion of neurotransmitter receptors in the plasma membrane of neurons is restricted by scaffolding proteins, which ensure the immobilization of receptors at the synaptic site (Choquet and Triller, 2013). Here, we present a novel model system that allowed us to investigate the interaction between scaffolding protein gephyrin and inhibitory GlyRs in their native oligomeric state of a pentamer. Based on our functional, structural, and cellular studies of GlyR-lCD-gephyrin interactions we propose a novel concept for a gephyrin scaffold-induced receptor oligomerization at the inhibitory synapse.

Introduction of α- and βICD did not affect the structural integrity of LS and enabled the formation of both LS-αICD and LS-βICD homopentamers as well as heteropentameric LS-αβICD. While the isolated LS-αICD showed a reduced folding stability as compared to LS^wt^ or LS-βICD, LS-αβICD was found to have the highest stability, which was attributed to additional interactions arising between GlyR α- and βICDs within the heteropentamer. Notably, full-length GlyRs comprising only β-subunits have not been described to build functional receptors, which suggests specific mechanisms, preventing GlyR β self-oligomerization. The formation of pentameric LS-βICD, shown in this study, indicates that ICDs alone could not hamper the interaction between individual LS-βICD entities and supports that rather the extracellular domain (ECD) and/or transmembrane domains (TM) prohibit the generation of GlyR β-homopentamers (Kuhse et al., 1993). On the other hand, the fact that LS-αβICD formed a stable and stoichiometrically defined complex, suggests that ICDs contribute to the subunit composition of GlyR. The herein determined assembly of two LS-αICD and three LS-βICD subunits within the pentameric LS-αβICD chimera is in line with the results of two independent previous studies on GlyR assembly (Grudzinska et al., 2005, Yang et al., 2012).

With our SAXS analyses we obtained first-in-class structural data on GlyR-ICDs providing a molecular view of the ICD interplay within the pentameric assembly. The SAXS models of LS-βICD and LS-αβICD support the dynamic nature of the GlyR-ICDs in GlyRs, which explains the lack of high-resolution structures of any pLGIC ICD in X-ray crystallography and cryo-electron microscopy studies (Du et al., 2015, Huang et al., 2015, Kumar et al., 2020). The obtained P1 and P5 models of LS-ICDs chimeras indicate the existence of multiple asymmetrical and symmetrical states, where GlyR-ICDs are extended to different degree from the “closed” conformation to the “open” one. Although the structures of both, LS-βICD and LS-αβICD, were highly similar, subtle but specific differences in the dimensionless Kratky plot suggest the formation of more compact assemblies within the heteromeric LS-αβICD. This finding is mirrored by the increased stability of LS-αβICD and dictates more efficient domain interactions between αICD and βICD in comparison to α/α and β/β interfaces. A similar structural anisotropy has been described for nAchRs, another member of the pLGIC receptor family (Miyazawa et al., 2003). The electron microscopy study of membrane-inserted nAchRs revealed in some receptors a vestibule structure of the ICD, suggesting a tight association of ICDs into a compact structure. On the contrary, the ICDs of the majority of the other nAchRs were not visible and appeared to lack any intracellular structure. These findings are in accordance with the structural variability observed for the GlyR-ICDs in this study and highlight a possible vital requirement of high ICD flexibility for GlyR interaction with postsynaptic membrane or other elements of the PSD (Ferraro and Cascio, 2018).

ITC studies showed that LS-βICD exhibited a twice as high saturation accompanied by a slightly decreased affinity (1.12 μM) towards GephE as compared to LS-αβICD (0.68 μM), suggesting steric interferences for the binding of the second GephE dimer within the ICDs pentameric assembly. In case of trimeric gephyrin, the affinity of interaction with LS-αβICD was even higher (0.49 μM) and in case of LS-βICD increased even further (0.26 μM). Binding stoichiometry between LS-αβICD was consistent with the binding of one gephyrin trimer, while the saturation of 0.93 reflects a mixture of one or two gephyrin trimers bound to LS-βICD. Finally, the interaction of both LS-βICD and LS-αβICD with gephyrin were characterized by an exothermic interaction with negative enthalpic-terms, being similar to the previously reported gephyrin-binding of GlyR-βICD peptides (Schrader et al., 2004, Maric et al., 2011, Specht et al., 2011, Herweg and Schwarz, 2012).

SEC studies revealed the formation of large complexes between gephyrin and LS-βICD- and LS-αβICD-specific masses. Equimolar interaction revealed complexes of similar sizes (approx. 2,000 kDa), corresponding to three gephyrin trimers and three LS pentamers (Figure 5). Further increase of gephyrin allowed the occupancy of additional binding sites on LS-variants leading to the binding of 1-3 additional gephyrin trimers with near proportional growth in size of the resulting complexes. While LS-βICD complexes represent an artificial accumulation of binding sites, their increased mass demonstrates the consistency in gephyrin-dependent formation of defined high-molecular aggregates in which gephyrin is able to link various pentamers.

In reverse, LS excess caused a reduction in complex size (1,570-1,710 kDa) representing more polydisperse species. This reciprocal relationship might indicate a key mechanism for ‘microcluster’ formation, which relies strongly on local changes of gephyrin or receptor concentrations at the PSD (Specht et al., 2020). All in all, our ITC and SEC data conclusively supports the view that a complex between three gephyrin trimer and three LS-αβICD pentamers is the most stable configuration, which correlates well with data received from EM-tomography experiments from native inhibitory PSDs (Liu et al., 2020).

The formation of high-molecular complexes between LS variants and gephyrin was further verified on cellular level by the accumulation of LS-βICD at synaptic sites in hippocampal neurons. Over-expression of LS-βICD increased the amount of gephyrin molecules at synaptic sites supporting the hypothesis that LS-βICD is able to link additional gephyrin molecules to the PSD at GABAergic synapses. This result underlines the hypothesis, that intermolecular binding events have to take place at the synapse to form a cluster spanning network and, that βICDs fulfil a bridging function during the clustering process of GlyR and gephyrin. In addition, *in vivo* data indicate a reciprocal mechanism of the regulation of GlyR-gephyrin networks, where the recruitment of both, receptor and scaffold, mutually depend on one another. Such a mutual stabilization of postsynaptic receptor-scaffold networks at synapses has been reported for GABA_A_Rs and gephyrin (Bannai et al., 2009) and is in line with our findings. Similarly, nAchRs were also reported to be bridged by the networking protein rapsyn at the postsynaptic membrane (Zuber and Unwin, 2013).

Recent studies have advanced the concept of phase separation as the driving force behind synaptogenesis (Zeng et al., 2016, Zeng et al., 2018). In this manner, phase separation at inhibitory post synapses was found to depend on receptor-scaffold interaction with the proposal that GephE dimerization is contributing to a linkage of different receptors (Bai et al., 2020). By preserving the native GlyR-ICD conformation and an αβ-subunit stoichiometry, our LS chimeras provide an excellent tool to enable further in-depth investigation by elucidating i. e. the role of alpha-ICD for the phase separation process.

In conclusion, the here discovered molecular mechanism underlying postsynaptic gephyrin scaffold formation might represent a universal concept for synapse formation of all pGLICs. Using LS as a model system, we are now able to mimic a cluster formation outside the membrane *in vitro*. The availability of LS-ICD models and their application for structural studies is an important step in deciphering the molecular architecture of the postsynaptic gephyrin-GlyR scaffold. It remains to be elucidated to which extend the different ICD conformations identified by SAXS will be able to preferentially bind to gephyrin and which other cellular factors will regulate their conformation during synaptic plasticity. Our LS-model may be applied to analyze ICDs of any member of pLGICs knowing that similar to GlyR, also ICDs of GABA_A_Rs as well as nAchRs serve as a critical determinant for receptor function (Zuber and Unwin, 2013, Tyagarajan and Fritschy, 2014). The LS backbone is therefore a novel and ideal tool to access so far unknown features of pLGIC ICD structure and function.

## Material and Methods

### Expression constructs

*Saccharomyces cerevisiae* LS sequence was cloned from yeast cDNA. Nucleotide sequences of rat GlyR-α_1_-ICD and rat GlyR β-ICD (corresponding to amino acid residues 338-427 and 334-454, respectively) were first inserted between Gly138 and Ile139 of LS via fusion PCR. Further, the obtained LS-αICD and LS-βICD constructs, as well as LS^wt^ were cloned separately in pQE70 vector (Qiagen, Hilden, Germany) via SphI/BglII for expression in *E. coli* as C-terminally 6xHis-tagged recombinant proteins. For co-expression of LS-αICD and LS-βICD, LS-αICD was inserted via SmaI/EcoRI in the first multiple cloning site and LS-βICD – via BglII/XhoI in the second multiple cloning site of pETDuet1 vector (Qiagen), leading to the attachment of the C-terminal 6xHis-tag to LS-αICD. For expression in COS-7 cells and hippocampal neurons LS-αICD, LS-βICD and LS^wt^ were cloned via BglII and SmaI into pEGFP-N2 vector resulting in fusion of GFP-tag at the C-terminus. LS-βICD was additionally introduced into pcDNA myc/His3.1 vector (Invitrogen, Carlsbad, California) and applied as C-terminally 6xHis- and myc-tagged construct for co-transfection with LS-αICD-GFP. Interaction studies were performed with recombinant full-length rat gephyrin splice variant rC4 cloned into pQE80 (Belaidi and Schwarz 2012). For expression in eukaryotic cells, gephyrin rC4 was inserted into pmCherry-C3 vector via XhoI/HindIII, leading to attachment of mCherry at the N-terminus.

### Expression and purification of recombinant proteins in *E. coli*

Gephyrin was expressed and purified according to Dejanovic et al. 2015 with additional washing steps during Ni-NTA chromatrography (45 and 60 mM imidazole). LS variants were expressed in *E. coli* BL21(DE3) for 3-5 h at 30-37 °C (LS^wt^, LS-βICD and LS-αβICD) and for 16 h at 18 °C (LS-αICD) upon induction with 100 μM isopropyl-β-D-thiogalactopyranosid at OD_600 =_ 0.6. Bacteria were harvested by centrifugation, resuspended in lysis buffer containing 50 mM sodium phosphate buffer pH 7.0-8.0, 300 mM NaCl, 10 mM imidazole, 5 mM β-mercaptoethanol and protease inhibitors (cOmplete, Roche, Basel, Switzerland) and lysed mechanically using EmulsiFlex (Avestin, Ottawa, Canada) and ultrasound for 1 min at 45 %. Cytosolic extract was separated from cell debris by centrifugation and purified via Ni-NTA chromatography. Ni-NTA matrix was washed in consecutive steps with lysis buffer containing either 40 mM and 100 mM imidazole (LS^wt^, LS-αICD and LS-βICD) or 25 mM and 40 mM (LS-αβICD) and eluted with lysis buffer containing 500 mM imidazole. Gephyrin and LS variants were further purified via size exclusion chromatography (Superdex 16/600 prep grade, GE Healthcare, Chicago, Illinois) in ITC-buffer. All purification steps were performed at 4°C.

### Analytical size exclusion chromatography

For determination of molecular masses purified LS variants were separated in analytical size exclusion chromatography (SEC) column Superdex 200 10/300 (GE Healthcare). Superose 6 increase 10/300 (GE Healthcare) was utilized for analysis of gephyrin and LS variants complexes. Thereby, 4 nmol, 8 nmol or 12 nmol gephyrin was mixed with 4 nmol (8 nmol or 12 nmol) of either LS-βICD, LS-αICD or LS-αβICD in 100 μl total sample volume. All SEC experiments were performed at a flow rate of 0.5 ml/min in a 25 mM Tris/HCl pH 7.5, 250 mM NaCl, 1 mM β-mercaptoethanol and 5% glycerol buffer. Molecular masses were calculated using gel filtration HMW calibration kit (GE Healthcare) (Figure S7a). To expand the calibration window, we included respiratory chain supercomplex I/III_2_/IV (1.7 MDa) prepared by solubilizing bovine heart mitochondria with 6 g/g digitonin and performing clear-native polyacrylamide electrophoreseis (Wittig et al., 2007) (Figure S7b). The supercomplex was eluted from the gel band in 0.1% digitonin, 50 mM NaCl, 1 mM EDTA, 2 mM imidazole pH 7.0 by diffusion.

### UV-Vis spectroscopy

UV-Vis spectra of purified LS variants were collected using GENESYS 10S spectrophotometer (ThermoFischer Scientific, Waltham, Massachusetts). Final protein concentration was adjusted to 40 μM LS^wt^, LS-αICD 40 μM, 44 μM LS-βICD and 40 μM LS-αβICD in the buffer containing 50 mM Tris pH 7.5, 350 mM NaCl and 15 μM β-mercaptoethanol

### Densitometric determination of band intensity

Densitometric determination of the intensity of protein bands of LS-αβICD was performed via a Coomassie-stained SDS-gel (12%) Therefore, increasing quantities (0.125 – 2 μg) of purified LS-αICD and LS-βICD were applied to the SDS-gel and were utilized as standard calibration curves. Either LS-αβICD or LS-αICD/LS-βICD mixed sample were applied onto the same gel in increasing manner (0.25 – 2 μg) to ensure that bands intensities were in the linear range of the respective calibration curve. All gels were recorded with a Chemidoc MP Imaging system (BioRad) and analyzed using Image LAB 5.2.1. software.

### Circular dichroism spectroscopy

Far-UV spectra were recorded with J-715 CD spectropolarimeter (Jasco, Gross-Umstadt, Germany) at 20 °C in the range of 190-250 nm using quartz cuvette with 0.1 cm path length. Final protein concentration was adjusted to 0.2 to 0.3 mg/ml in 50 mM Na-phosphate buffer pH 5.0,100 mM NaCl. Buffer baseline was recorded separately and subtracted from each sample spectrum. Obtained ellipticity (θ) was normalized to protein concentration in mg/ml (c), molecular mass in Da (Mr), number of amino acids (n) and path length of cuvette (l) using formula formula: [θ] = θ * Mr / 10 * (n-1) * c * l. Thermal unfolding of purified proteins with concentration of 0.15 to 0.25 mg/ml was performed in buffer containing 20 mM sodium phosphate pH 6.8 and 300 mM NaCl. Ellipticity was measured continuously at 220 nm in a sealed cuvette with a heating speed of 1°C/min from 10 °C to 90 °C and data pitch of 0.2 nm. Melting curves were fitted using Jasco Spectra Manager 2 software.

### Analytical ultracentrifugation (AUC)

All experiments were performed with a Beckman Optima XL-A instrument equipped with absorbance optics at a temperature of 4°C. For sedimentation equilibrium (SE) centrifugation 120 μl samples with 16 μM protein concentration were analyzed in standard aluminum two-channel centerpieces (1.2 cm path) after attaining equilibrium at different speeds using Tris/HCl pH 7.5, 250 mM NaCl, 1 mM TCEP and 5% glycerol as solvent. The distribution of the proteins was monitored at 280 and 285 nm. The 8-hole rotor AnTi-50 as well as sample cells had been pre-cooled. Scans were recorded using 0.002 cm point spacing and averaging 10 readings at each point. Data collected at different speeds were fitted globally by the global-fit function provided in UltraScan II vs 9.9.(Demeler, 2005, Demeler, 2018) Based on the fit result, finally, a Monte Carlo statistics with 10,000 iterations was done.

Partial specific volumes of the proteins, as well as density and viscosity of the buffer were calculated with the aid of SEDNTERP(Laue et al., 1992, Laue, 2018).

### Differential scanning fluorimetry (DSF)

Purified proteins with final concentration of 0.5 mg/ml in 20 mM sodium phosphate pH 6.8 and 300 mM NaCl buffer were supplemented with SYPRO Orange (ThermoFisher Scientific) accordingly to manufacture’s specifications and heated up in sealed qPCR stripes (Bio-Rad, Hercules, California) from 20°C to 95 °C in C1000 Thermal Cycler CFX96 Real-time System (Bio-Rad). T_m_ was determined as a local minimum of the first derivative of the raw fluorescence as function of temperature with software Bio-Rad CFX Manager 3.1.

### Isothermal Titration Calorimetry

Experiments with gephyrin trimer or GephE titrated into LS variants were performed by using MicroCal Auto-ITC200 (Malvern, Malvern, United Kingdom) at 25 °C in a 25 mM Tris/HCl pH 7.5, 250 mM, 5 mM β-ME and 5 % glycerol buffer. The injection volume was set to 1-2 μl with a spacing of 150 s between injections and an initial delay of 60 sec. The reference power was set to 5 μCal/sec and the stirring speed to 1000 rpm. Gephyrin concentration in the syringe was set to 170 -220 μM, GephE concentrations were set to 200-250 μM and LS variant concentration in the sample cell to 15 – 25 μM.

All experiments were performed using proteins from a minimum of two individually prepared purifications for each interaction partner. ITC isotherms were fitted using the software Origin 7. Errors represent standard error of the mean (SEM), calculated from all measurements.

### Cell culture, transfection and immunostaining

Murine primary hippocampal neurons were prepared from day 17.5 embryos and cultured in Neurobasal medium (Gibco, ThermoFischer Scientific) supplemented with glutamine, B27 and N2 (Gibco, ThermoFischer Scientific). Neurons were transfected with Lipofectamine 2000 (Invitrogen) at DIV 9 according to the manufacturer’s protocol and immunostained at DIV 11. COS-7 cells were cultured in DMEM (GE Healthcare) supplemented with FCS (PAN-Biotech, Aidenbach, Germany) and glutamine (Gibco, ThermoFischer Scientific). COS-7 cells were transfected using polyethylenimin (PEI) and stained after 24 h.

For immunostaining, both neurons and COS-7 cells were fixated with 4% PFA, the reaction was stopped with NH_4_Cl. Cells were blocked in PBS with 1 % BSA, 10 % goat serum and 0.2 % Triton X-100. Primary antibodies anti-gephyrin (1:50, cell culture supernatant, clone 3B11, Synaptic Systems) and anti-GABA_A_R subunit α2 (1:50, Synaptic Systems, Goettingen, Germany) and secondary antibodies Alexa Fluor 568, 488 (1:500, ThermoFisher Scientific) and Pacific Blue (1:500, ThermoFisher Scientific) were diluted in blocking buffer. Nuclei were stained with DAPI (1:1000 in PBS, AppliChem, Darmstadt, Germany) and coverslips were mounted on Mowiol 4-88 (Calbiochem, Merck Millipore, Billerica, Massachussetts) with the addition of 2,5% 1,4-Diazobicyclo[2.2.2]octan (Dabco, Merck Millipore), adjusted to pH 9.0.

### Image acquisition and analysis

Microscopic images were acquired with an inverted laser scanning microscope (Eclipse-Ti, Nikon A1, Nikon, Shinagawa, Japan) with a resolution of 1024 × 1024 pixels (4.823 pixels / μm). Pictures were recorded with the EZ-C1 software (version 3.60, Nikon). Neuronal images represent a maximum intensity projection of a Z-stack of three optical sections with 0.45 μm steps. Images of COS-7 cells are average-pictures of 4 images. Images were processed with the software ImageJ.

For analysis, clusters of at least 15 neurons per construct were quantified from three sets of neurons. Settings were determined and stayed constant during each set to allow comparable results. Dendritic clusters in two 20 × 5 μm regions of interest (ROIs) per neuron were counted, their area and fluorescent intensity measured with the NIS-elements software (Nikon). Shown values represent averages. Statistical analysis was performed using Microsoft Excel. Significance of mean values were calculated with student’s t-test, errors represent standard error of the mean (SEM).

### Small-angle X-ray scattering (SAXS)

Small-angle X-ray scattering data were recorded at the BioSAXS BM29 beamline (Pernot et al., 2013) at the European Synchrotron and Radiation Facility (ESRF) in Grenoble, France. To insure the monodispersity of samples all proteins were subjected to SEC directly before collecting of scattering data on a Superdex 200 10/300 GL (GE Healthcare) equilibrated with 20 mM Tris pH 7.5, 300 mM NaCl and 15 mM beta-mercaptoethanol (Figure S7). Fractions corresponding to pentamers were collected, concentrated with Centricon (Merck Millipore) devices and either measured in static mode (LS^wt^) using sample changer (Round et al., 2015) or reapplied on the SEC column coupled to the flow-through SAXS capillary (LS-βICD and LS-αβICD) (Figure S7d and e). Scattering of three different LS^wt^ concentrations (6.5, 3.4 and 1.2 mg/ml) was analyzed and merged with PRIMUS (Konarev et al., 2003) of the ATSAS suite (Petoukhov et al., 2012). The scattering of SEC buffer was measured before and after each LS^wt^ sample and subtracted automatically with the beamline control software (Pernot et al., 2013). Following the buffer subtraction, the scattering curve was submitted to Guinier analysis in PRIMUS, whereby the signal intensity at small scattering angles was used to calculate radius of gyration (R_g_) and forward scattering I(0). Computation of inter-atomic distance function *P*(*r*) was performed with GNOM (Svergun, 1992). In the online SEC mode the scattering intensities and absorption at 280 nm were collected continuously over the SEC-column volume (Figure S7c-e). The frames corresponding to SEC-baseline prior protein peaks were inspected, merged and subsequently used for buffer subtraction. The frames corresponding to LS-βICD or LS-αβICD with a consistent R_g_ (Figure S7d and e) were merged and used for computation of *P*(*r*) function and subsequent structural modelling. *Ab initio* modelling with five-fold symmetry and without symmetry constrains was performed with DAMMIF (Franke and Svergun, 2009) and GASBOR (Svergun et al., 2001) generating 20 and 10 independent models, respectively. DAMMIF models were subsequently averaged with DAMAVER (Volkov and Svergun, 2003). Evaluation of the experimental scattering data for LS^wt^ was carried out with CRYSOL (Svergun et al., 1995) using crystal structure of LS from *S. cerevisiae* (PDB 1EJB). Molecular masses of LS variants were calculated directly from SAXS scattering applying SAXS MoW for LS^wt^ (Fischer et al., 2010) and SCÅTTER for LS-βICD and LS-αβICD (Rambo and Tainer, 2013).

## Acknowledgements

We thank Manuel Martinez-Osuna, Joana Stegemann and Monika Laurin (University of Cologne, Germany) for the technical support. Financial support by the German Science Foundation (DFG, SFB 635) is gratefully acknowledged. We thank the European Synchrotron Radiation Facility (Grenoble, France) for provision of synchrotron radiation facilities and we would like to thank Martha Brennich for assistance in using beamline BioSAXS BM29. We acknowledge European Molecular Biology Laboratory (Grenoble, France) for provision of lab facilities. The authors declare no financial competing interest.

## Supplementary

**Figure S1.**
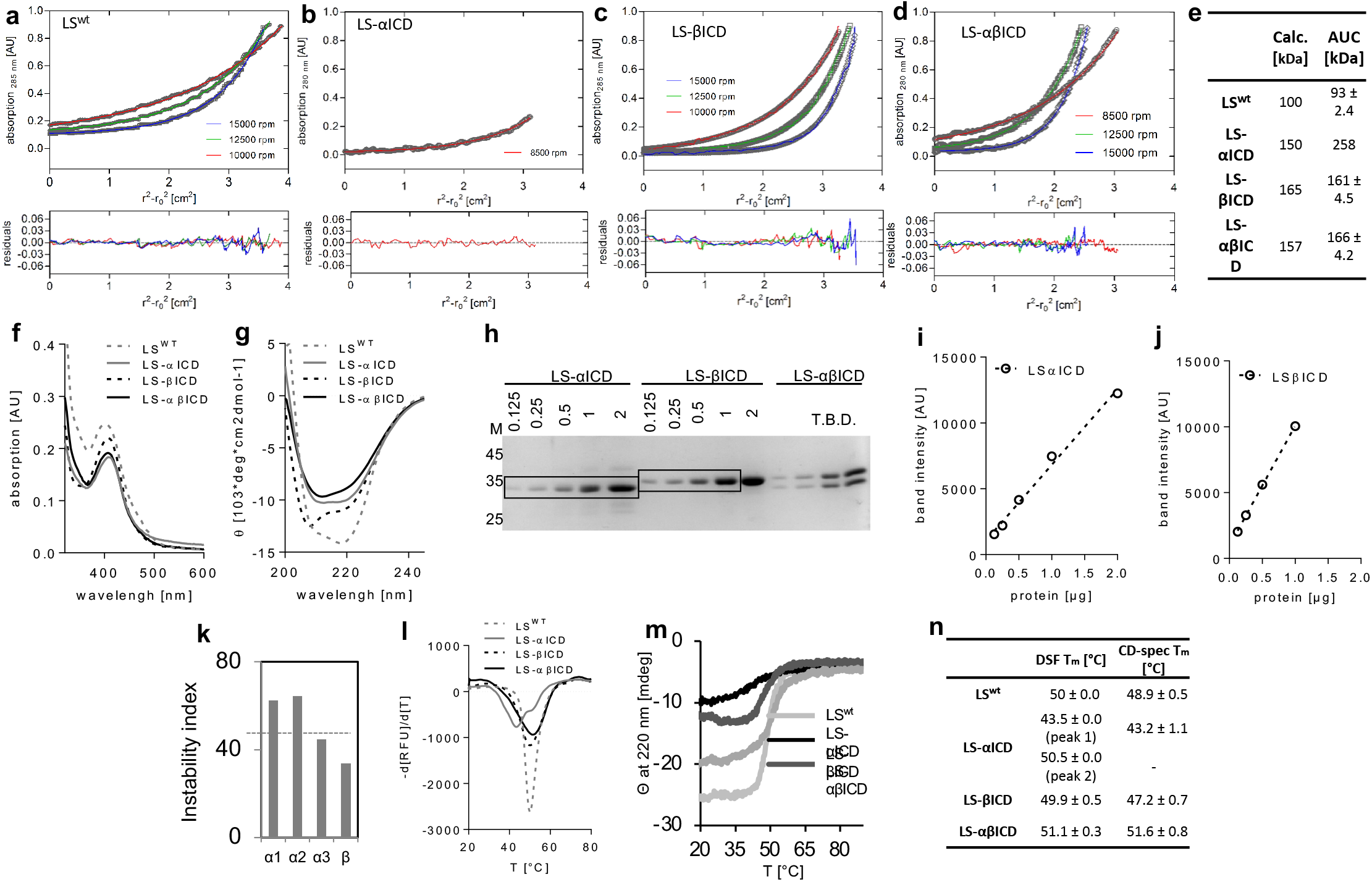

**Figure S2.**
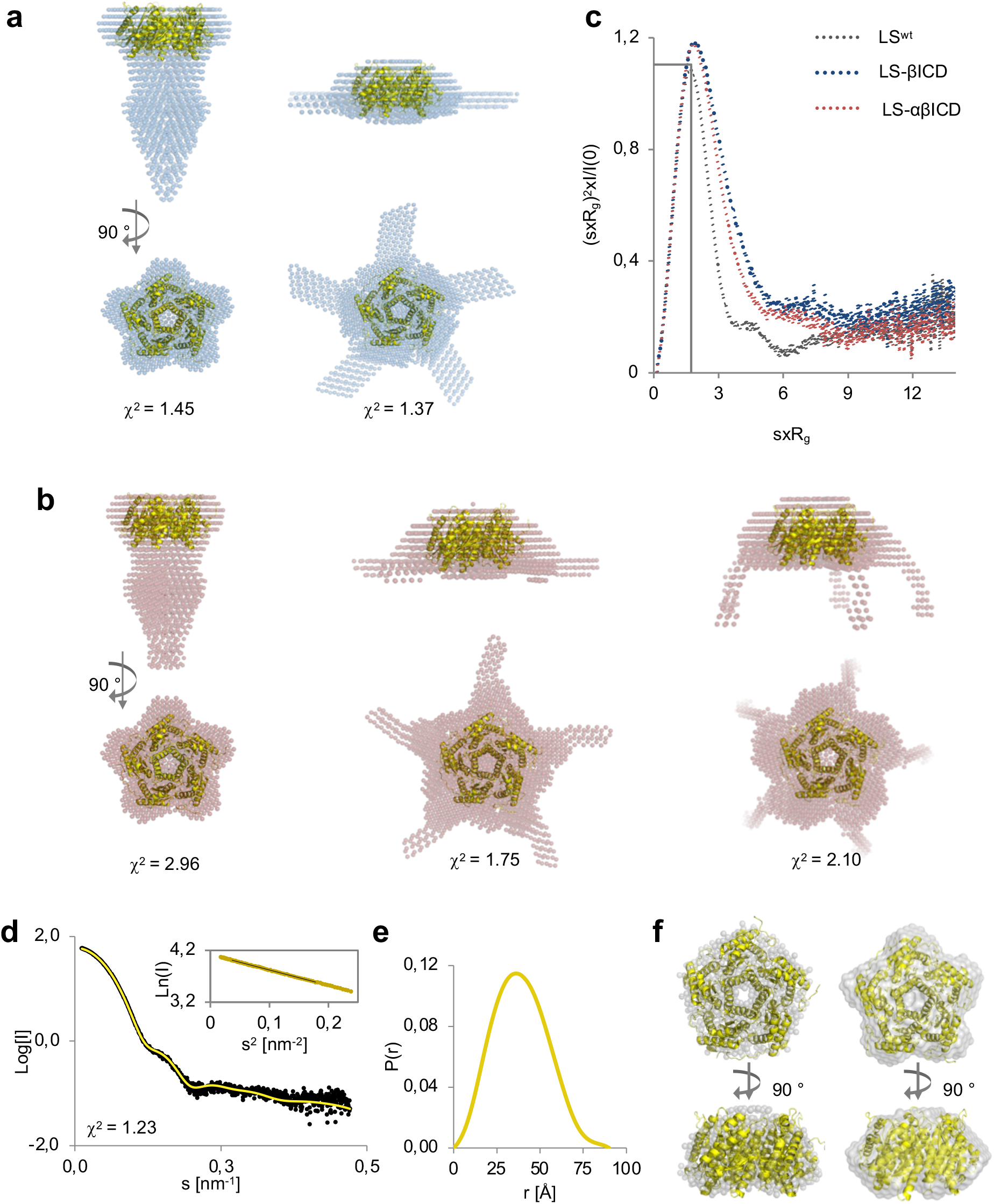

**Figure S3.**
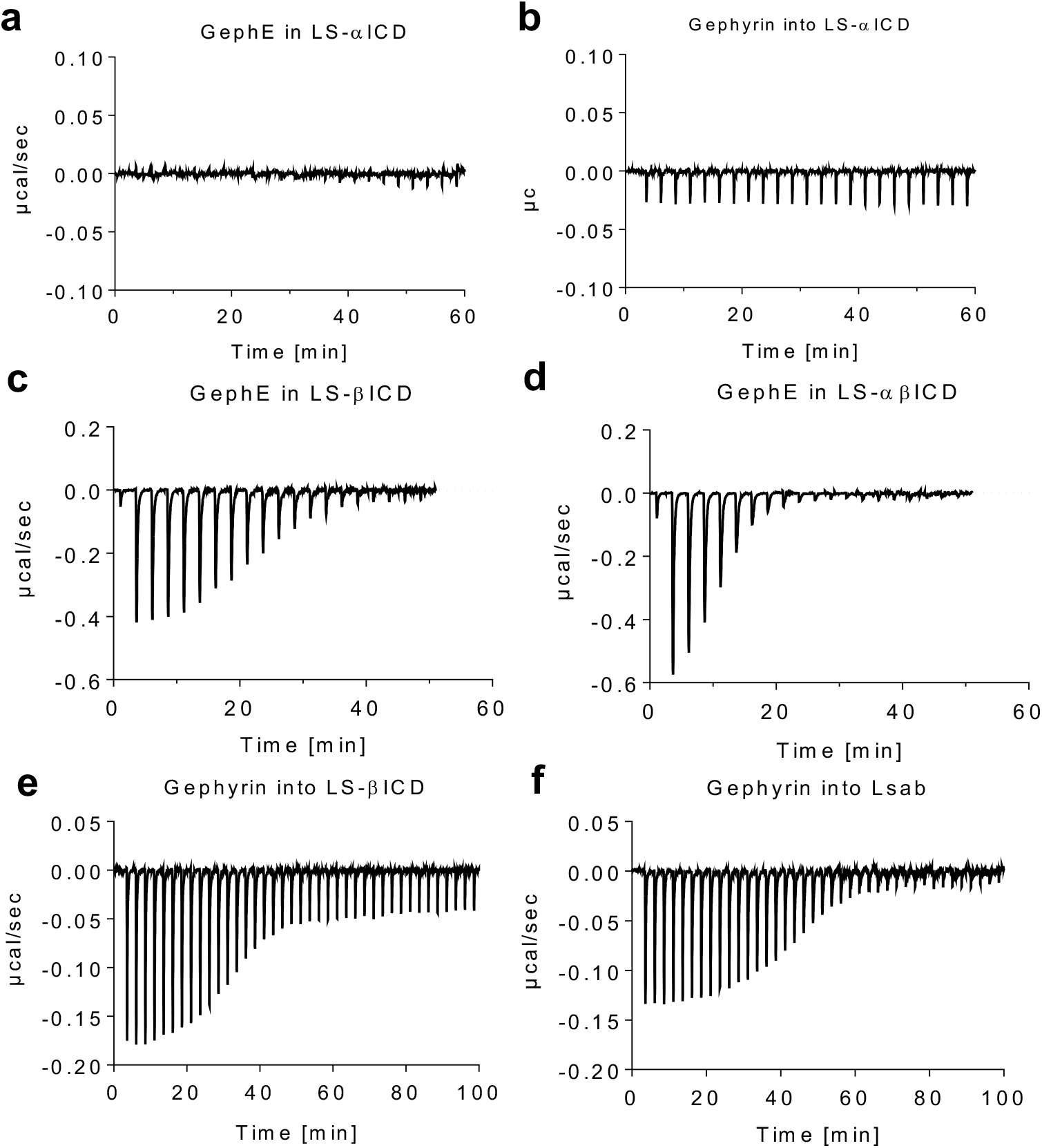

**Figure S4.**
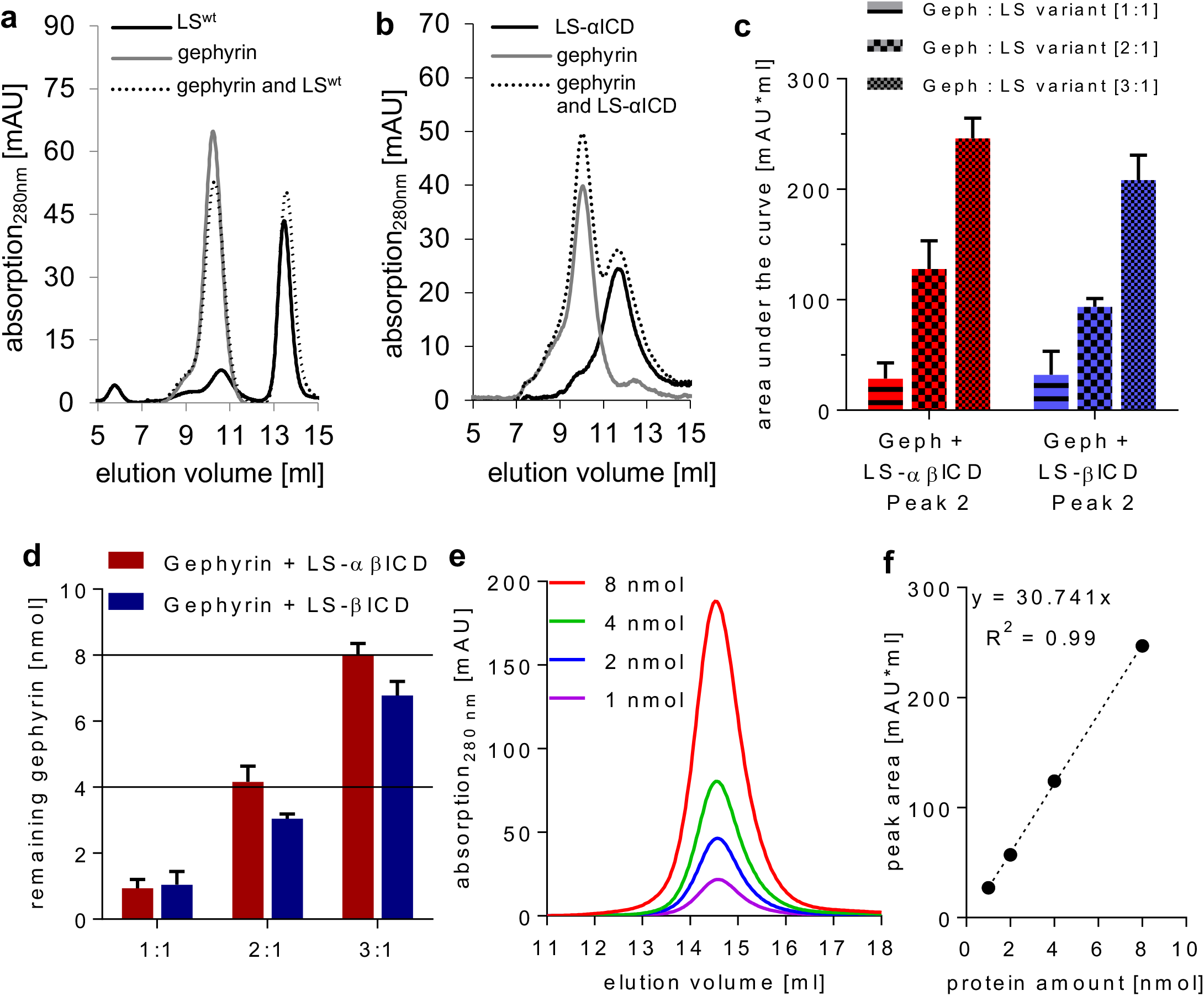

**Figure S5.**
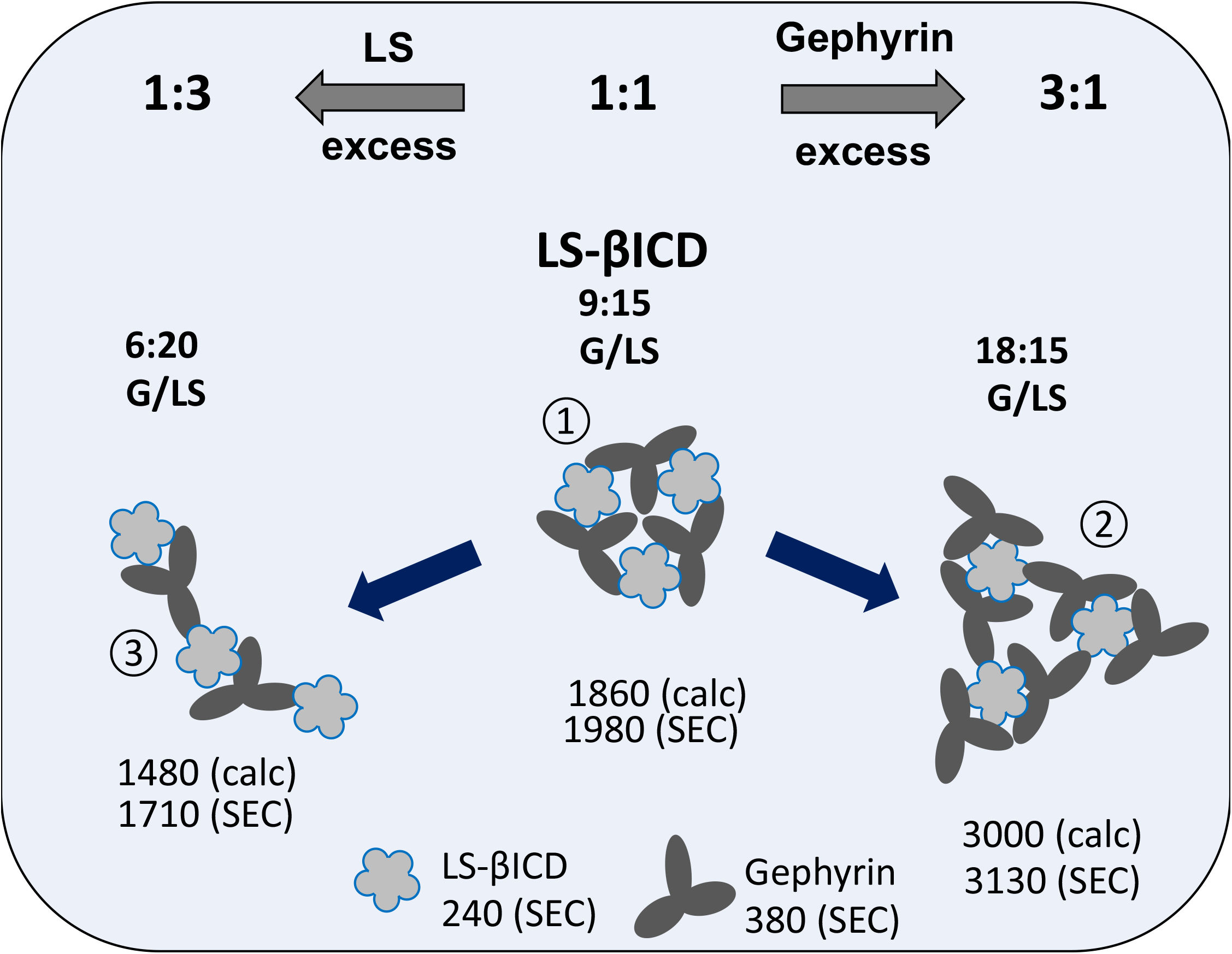

**Figure S6.**
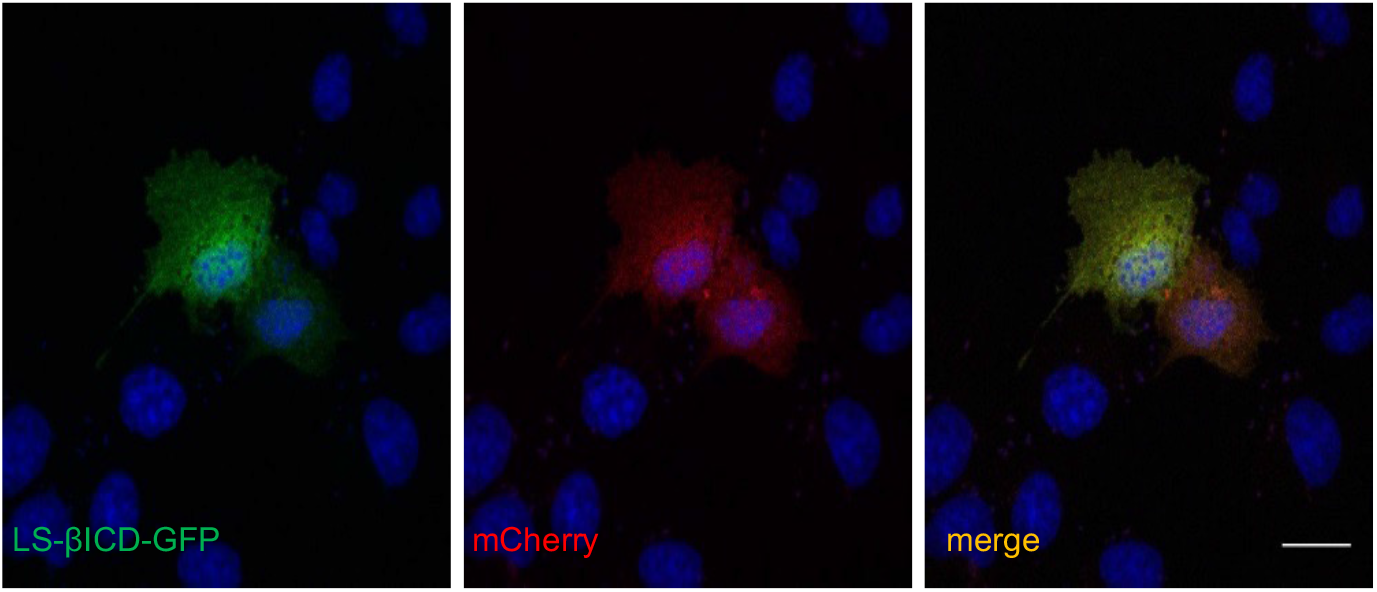

## Supplementary figure methods

**Figure S7.**
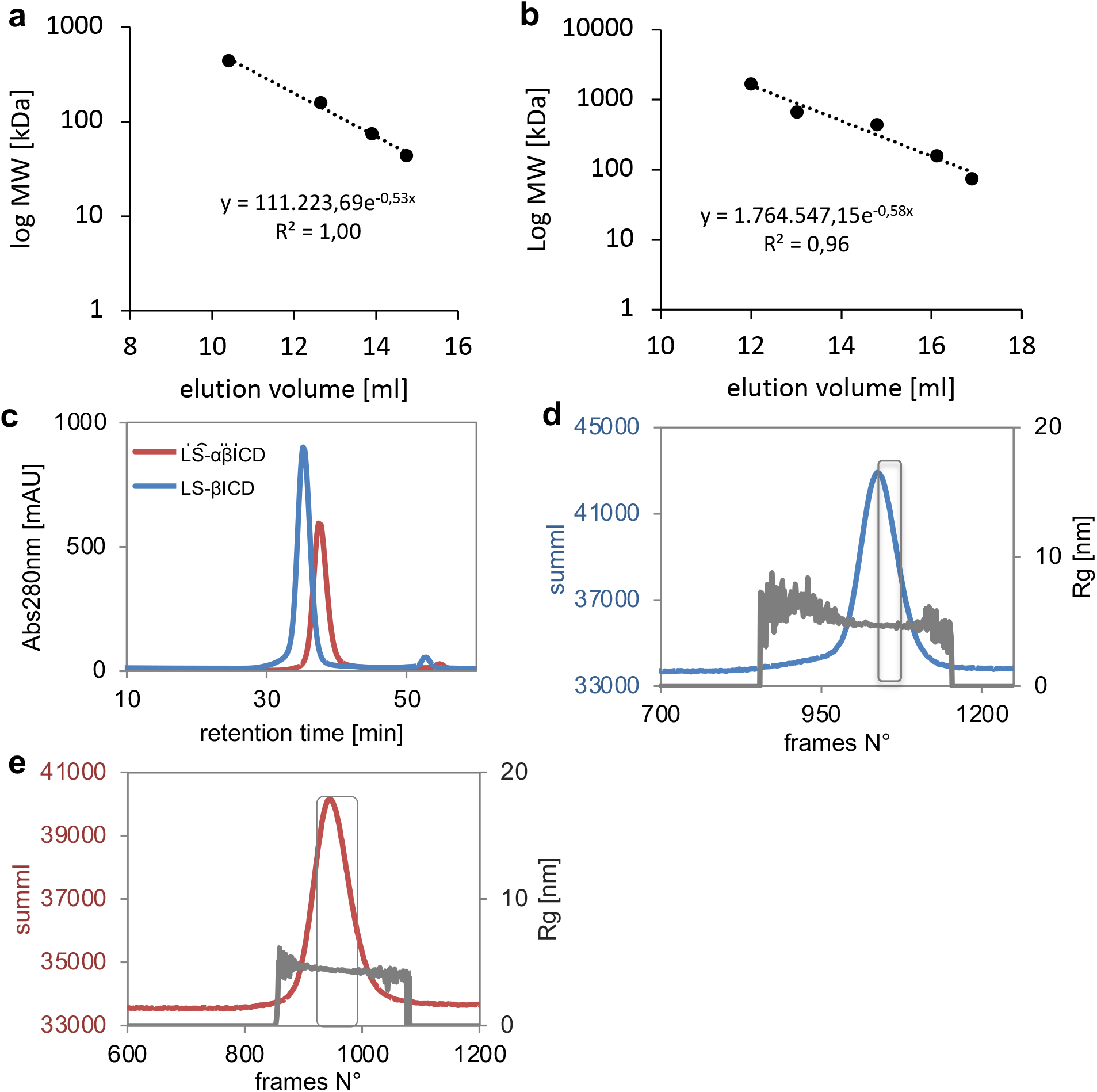

