## Supplementary Figures for "Pentameric assembly of glycine receptor intracellular domains provides insights into gephyrin clustering"

**Supplemental Information**

Arthur Macha[1,#], Nora Grünewald[1,#], Nastassia Havarushka[1,#], Nele Burdina[1],
Christine Tölzer[1], Luitgard Nagel-Steger[2], Yvonne Merkler[1], Karsten Niefind[1],
Guenter Schwarz[1,2]*

[1] Institute of Biochemistry, Department of Chemistry, University of Cologne, 50674 Cologne, Germany

[2] IBI-7, Structural Biochemistry, Forschungszentrum Jülich, 52425 Jülich & Institut für

Physikalische Biologie, Heinrich-Heine-Universität Düsseldorf, 40225 Düsseldorf, Germany

[3] Center for Molecular Medicine Cologne, University of Cologne, 50674 Cologne, Germany

[#]contributed equally

### Supplemental Results

#### Biochemical characterization of LS variants

Application of LS as an ICD-carrying platform relies on correct oligomerization and folding of LS monomers following the introduction of the respective ICDs. It is known, that upon expression in *E. coli*, yeast LS is isolated as a yellow-colored protein due to co-purified riboflavin (Woycechowsky et al., 2006). As the active site of LS is situated at the interface of the adjacent subunits within the pentamer, one can assume that disruption of the inter-subunit interface would lead to the loss of the binding pocket at the active site and subsequent colorless appearance of the obtained proteins. In agreement with a native fold of LS, all purified LS chimeric proteins showed yellow-colored appearance as judged by UV/vis spectroscopy (Figure S1f). Thus, the structural integrity of the pentameric backbone, necessary for the riboflavin binding, remained unaffected upon insertion of GlyR-ICDs.

The impact of the GlyR-ICD on the secondary structure of LS was further analyzed by CD-spectroscopy (Figure S1g). $LS^{wt}$ showed in the far-UV range a high content of α-helices as represented by two negative maxima at 208 and 222 nm, which is in agreement with previously reported CD-spectra (Woycechowsky et al., 2006). The insertion of neither GlyR $\alpha_1$- nor GlyR β-ICD diminished the characteristic biphasic shape of the CD-profile, confirming preservation of the $LS^{wt}$ secondary structure. The observed difference in the curve progression between $LS^{wt}$ and the LS-ICD variants suggested a specific folding of the corresponding ICDs. Changes in the intensity of major peaks at 208 nm and 222 nm might indicate an impact of LS-αICD on the structure of LS-βICD in the heteromeric LS-αβICD protein (Figure S1g).

The influence of the subunit composition on the overall stability of LS assemblies was investigated. Analysis of predicted intrinsic stability of GlyR subunit sequences for their content of destabilizing dipeptides (Gasteiger et al., 2003, Guruprasad et al., 1990) showed that, differently to GlyR β-ICD, all isoforms of the GlyR α-ICD were predicted to be intrinsically unstable (Figure S1k). Therefore, thermostability of LS-ICD chimeras were examined by differential scanning fluorimetry (DSF) and CD-spectroscopy (Figure S1l-n). LS-βICD showed a melting temperature ($T_m$) of approximately 50°C, which was similar to that of $LS^{wt}$, indicating that β-ICD alone neither contributes to further stabilization of the complex, nor does it cause any destabilization on the chimeric variant. In contrast, unfolding of LS-αICD was observed at considerably lower temperatures as depicted by the reduction of the transition point in the CD-curve to 42.7 °C as well as by the appearance of an additional peak at 43 °C in the DSF analysis (Figure S1l-n). Notably, in LS-αβICD the presence of βICD compensated for the destabilizing action of αICD, as the thermal melting of LS-αβICD reached a $T_m$ of 51 °C, which indicates additional stabilizing interactions between αICD and βICD within the heteropentamer.

**Small angle X-ray scattering**

First, SAXS data of $LS^{wt}$ without attached GlyR-ICDs were collected (Table 1 and Figure S2d-f). The scattering curve of $LS^{wt}$ was linear within the Guinier region confirming sample homogeneity, which is required for reliable structural modelling (Fig S2d). The inter-molecular distance distribution function *P*(r) derived from the scattering curve, exhibited a symmetrical profile, as it is expected for compact and spherical particles (Figure S2e) (Svergun, 1992). The estimated molecular mass of 106 kDa was in agreement with the theoretical mass of the LS pentamer (99.5 kDa, Table 1).

*Ab initio* models of $LS^{wt}$ structure in solution were generated using DAMMIF and GASBOR. The best fit of the experimental data was achieved following the application of a 5-fold symmetry, depicturing the distinct conical shape of LS and the pore in the center of the LS-pentamer (Figure S2f). Upon omission of symmetry constrains, both DAMMIF and GASBOR yielded an oblate molecular envelop with the dimensions matching the model derived from the crystal structure of $LS^{wt}$ and a slightly irregular contour (Figure S2f). The-goodness-of-fit ($\chi^2$) between experimental and the theoretical scattering calculated from the crystal structure gave value of $\chi^2$ = 1.23 (Figure S2d), demonstrating that the conformation of $LS^{wt}$ in solution was highly alike to that of the crystal structure. Next, the scattering curves of LS-βICD and LS-αβICD were examined. $R_g$ values derived from the Guinier fit and from the $P(r)$ function were consistent for both proteins (Table 1).

Models of LS-αβICD and LS-βICD generated with DAMMIF (P5 symmetry), in contrast, were organized in several different clusters, where the ICDs were either separated from each other similarly to the GASBOR models, or were positioned together, building a prolonged rod stemming from the center of the LS (Figure S2a and b). LS-αβICD yielded more clusters than LS-βICD presumably illustrating higher anisotropy of LS-αβICD conformers due to the presence of the GlyR-ICDs derived from α-subunit.

The flexibility of GlyR-ICDs within LS-βICD and LS-αβICD was further addressed by applying dimensionless Kratky plot of the scattering data, which allows the comparison of particles independent of their concentration and mass. In the corresponding plot of $LS^{wt}$, the curve reached its maximum at 1.101 at $R_g \times s$=1.73 (Figure S2c), being characteristic for globular, compact proteins (Durand et al., 2010). Differently, the maxima of LS-βICD and LS-αβICD were shifted to higher values and the function became more extended with the increasing $R_g \times s$. Thus, the hypothesis was supported that a considerable degree of flexibility has been introduced into LS-βICD and LS-αβICD due to the presence of the ICDs (Receveur-Brechot and Durand, 2012). Notably, the scattering pattern of LS-βICD was slightly more extended in comparison to LS-αβICD. This might be indicative for more pronounced interactions between α- and β-ICDs within LS-αβICD in comparison to β-ICD/β-ICD interfaces within LS-βICD.

### Supplemental Figures

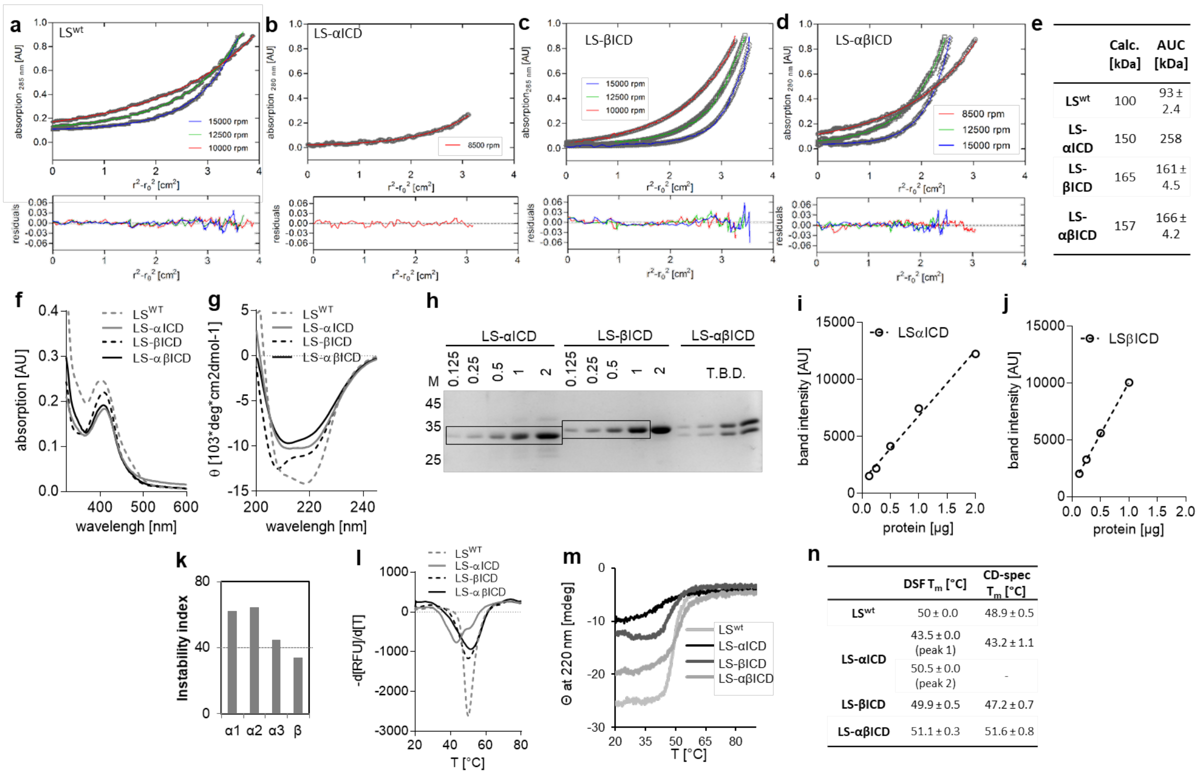


**Figure S1: Biochemical characterization of LS variants. (a-d)** MWs determination of LS-variants using sedimentation equilibrium experiments of (a) LS$^{wt}$ (b) LS-αICD (c) LS-βICD and (d) LS-αβICD at various velocities (indicated). The upper panels show concentration profiles recorded after the establishment of equilibrium between sedimentation and back diffusion and the calculated concentration distributions. Lower panels depict the respective residuals of the fits above. Determined MW for each LS variant with technical errors are indicated. Note for LS-αICD, at velocities higher than 8500 rpm no protein was detected in the AUC sample. **(e)** Summary of determined MWs via AUC for LS variants in comparison to the calculated MWs. **(f)** UV-Vis spectra of LS variants, which were purified as yellow-coloured solutions with broad absorption maxima at approximately 410 nm. **(g)** Secondary structures profiles of purified proteins obtained via CD-spectroscopy. **(h)** Representative SDS-PAGE of purified LS-αICD and LS-βICD applied in various total protein amounts [µg] in addition to purified LS-αβICD. Note, for LS-βICD the 2 µg band was not considered for the calibration curve, due to signal intensity exceeding the detection limit. **(i, j)** Resulting calibration curves of LS-αICD (i) and LS-βICD (j), used for the calculation of total protein amount of α and β subunit in LS-αβICD based on the determined band intensities in (h). **(k)** Predicted intrinsic stability of GlyR-ICDs derived from α1- , α2-, α3- and β-subunits. Proteins with instability index below 40 (dashed line) considered as stable (Gasteiger et al., 2003, Guruprasad et al., 1990). **(l)** Melting curves obtained during differential scanning fluorimetry (DSF). **(m)** CD-melting curves of LS$^{wt}$ and LS-variants. **(n)** Summary of melting temperatures ($T_m$) determined with DSF and CD-spectroscopy.

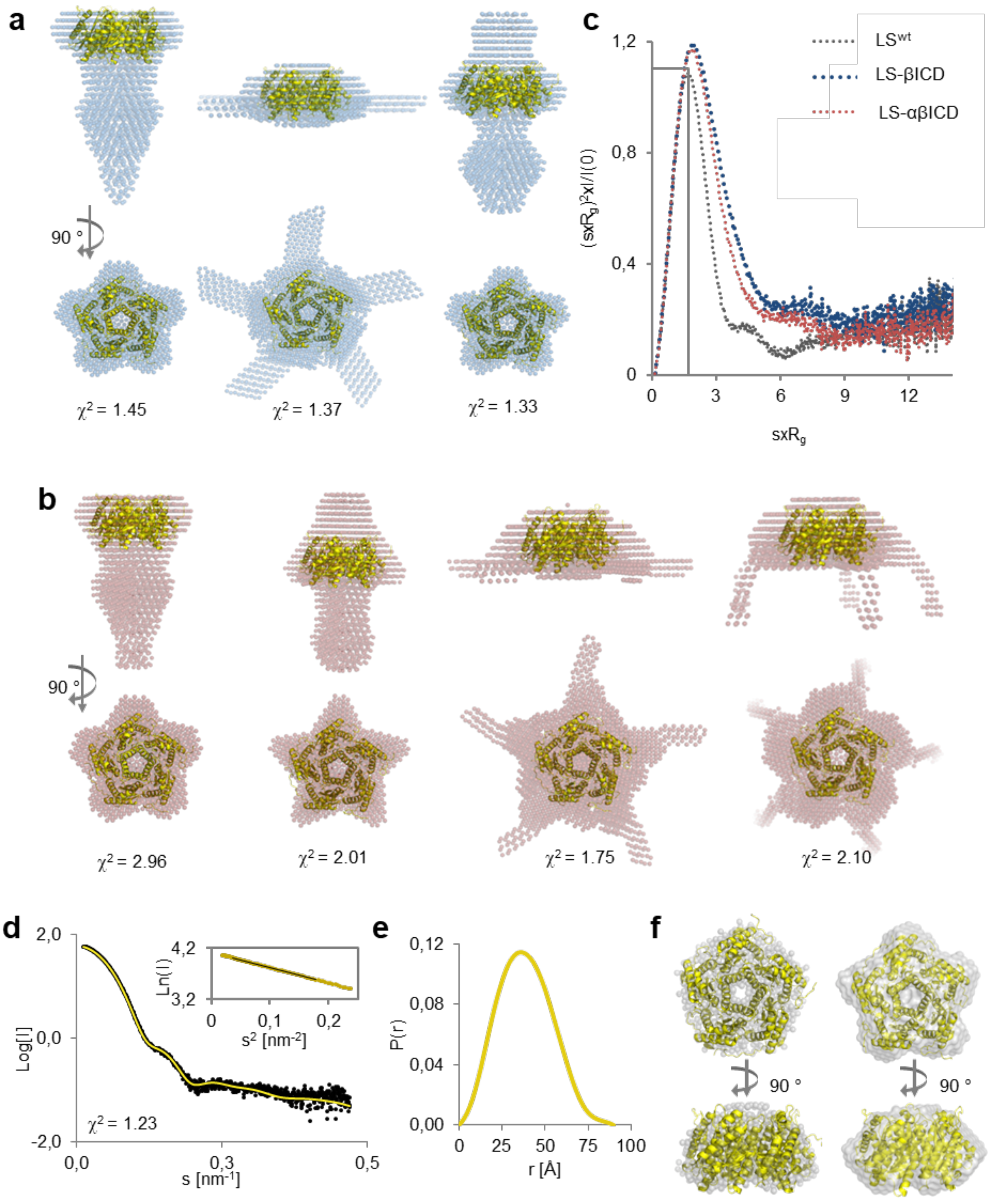


**Figure S2. Ab initio DAMMIF models of LS-βICD, LS-αβICD and LS$^{wt}$. (a-b)** Representative P5 DAMMIF models of each cluster obtained for LS-βICD (blue) and LS-αβICD (red) superimposed with LS crystal structure (yellow). Goodness-of-fit (χ2) values of each model are shown. **(c)** Dimensionless Kratky plot of LS variants in comparison to LS$^{wt}$. **(d)** CRYSOL fit of the experimental intensities (black dots) and intensities calculated from the LS crystal structure

(PDB 1EJB, yellow line). Insert: linear fit (black line) of the scattering curve within Guinier region (yellow). **(e)** P(r) function of the $LS^{wt}$ scattering curve. (f) Ab initio P5 models calculated with GASBOR (left) and DAMMIF (right) shown in grey and superimposed with LS crystal structure (yellow). Water molecules in the representative GASBOR model were omitted for clarity.

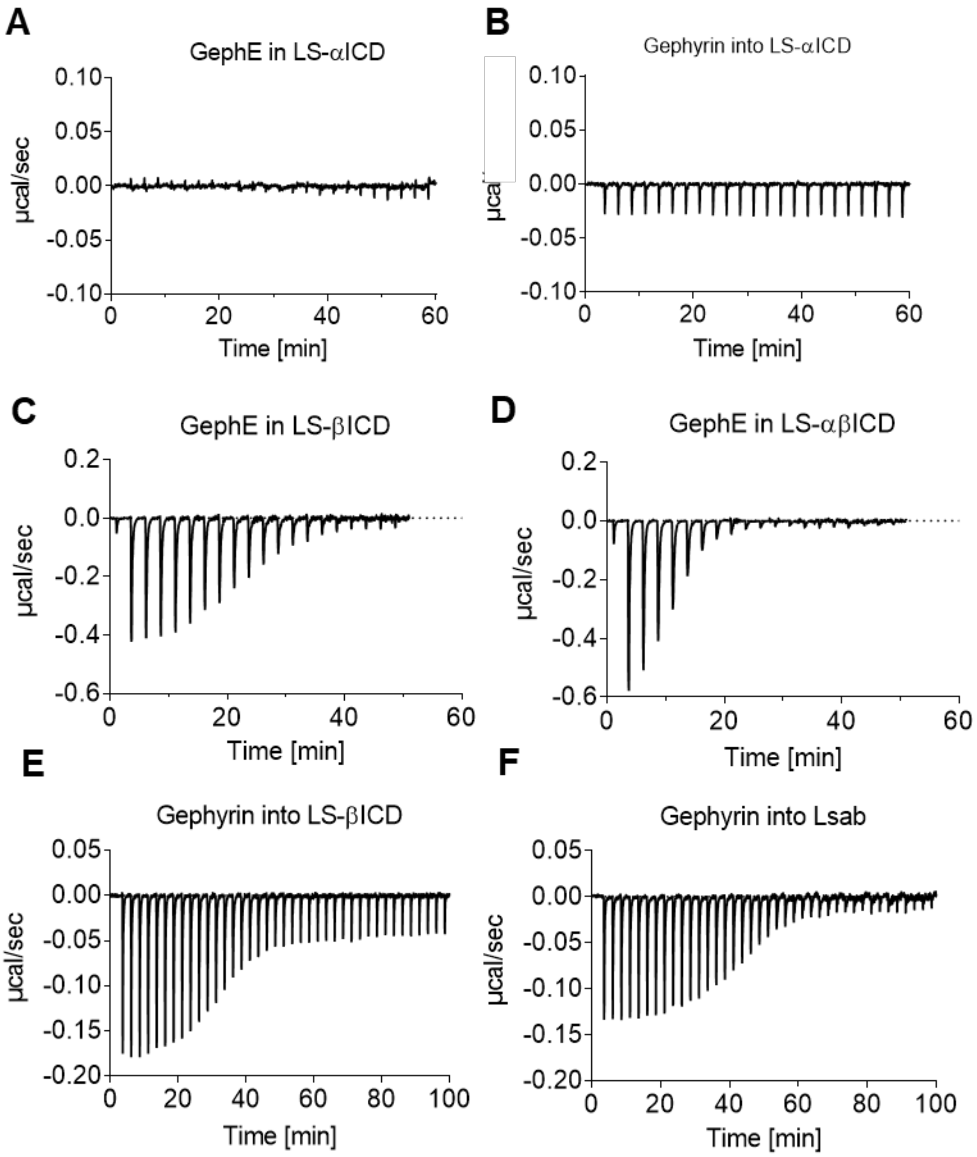


**Figure S3**. **Overview of representative ITC thermograms** of **(a, b)** Gephyrin variants into LS-αICD, **(c, d)** GephE into LS-variants and **(e, f)** gephyrin into LS-variants.

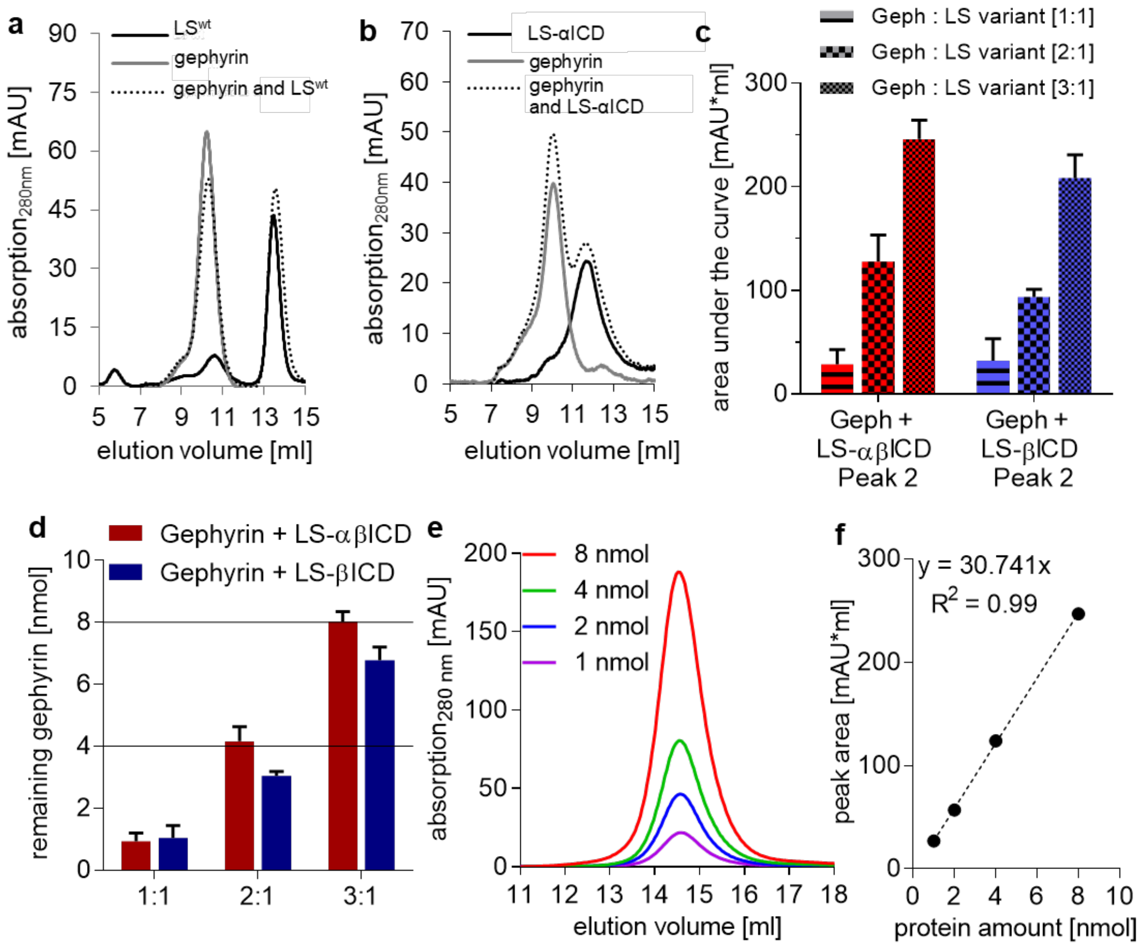


**Figure S4. Analysis of interaction between gephyrin and LS-variants via SEC**. **(a, b)** Gephyrin (4 nmol) was incubated with $LS^{wt}$ (4 nmol) (a) or LS-αICD (4 nmol) (b) prior to the application onto the analytical SEC column (dashed lines). All proteins were also separated individually (gephyrin: grey lines, LS variants: black lines). **(c)** Determined area under the curve of peak 2 of respective SEC experiments depicted in Figure 4a, c at equimolar protein amounts (1:1, horizontal stripes), two-fold (2:1, black squares) and three-fold (3:1, small squares) gephyrin excess. Data presented as mean ± SD of n = 3 independent experiments. **(d)** Calculated remaining total protein amounts [nmol] of gephyrin derived from peak areas in (c) by using protein calibration curve (f). **(e)** Representative SEC traces of purified gephyrin trimer applied in various amounts (1-8 nmol). **(f)** Calibration curve of the areas under the curve determined in (e) plotted against the correlating protein amount (black dots) with linear fit (dotted line). Function of the linear fit and goodness-of-fit ($R^2$) are indicated.

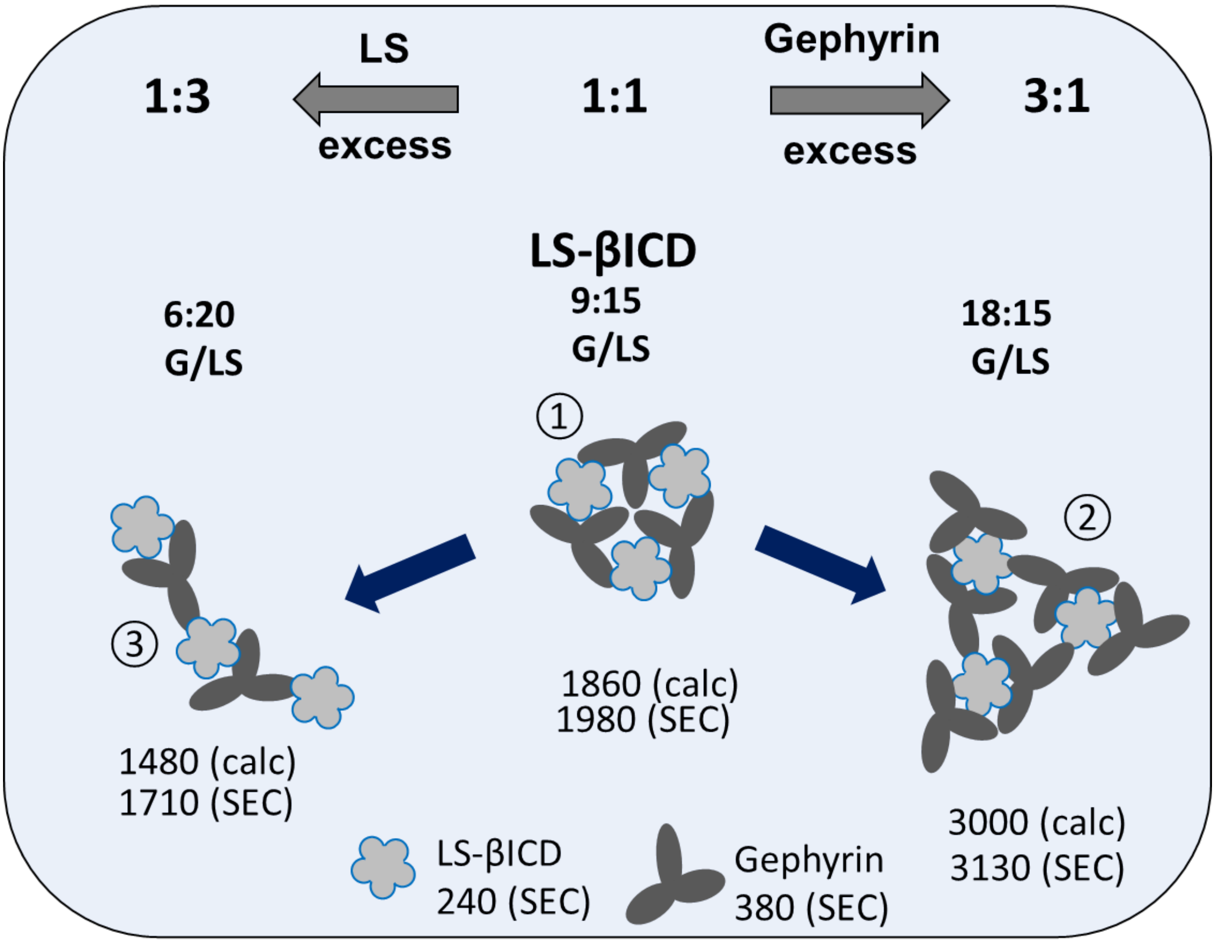


**Figure S5: Model of gephyrin-LS-βICD complex formation.** Depicted are the respective complexes formed at equimolar quantities (1:1) and with three-fold gephyrin (3:1) or LS (1:3) excess. In a 1:1 equimolar setup LS-βICD variant was saturated with the same amount of gephyrin molecules, resulting in a complex of three gephyrin trimer and three LS pentamers with MWs of 1980 kDa (1). Three-fold gephyrin excess led to an enlargement of the complex by three gephyrin trimer, increasing the MW to 3130 kDa (2). LS excess resulted in the formation of smaller complexes (1710 kDa) through the release of gephyrin molecules, probably due to the oversaturation of gephyrin binding sites (3).

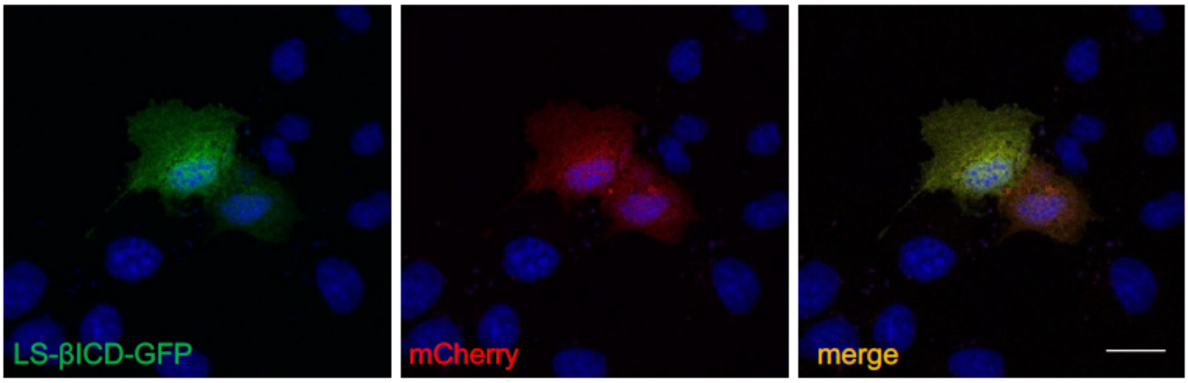


**Figure S6. Control transfection of LS-βICD-GFP and mCherry in COS-7 cells**. LS-βICD-GFP (green) and mCherry (red) were co-transfected in COS-7 cells (merge).

**Supplemental Figures for method section**

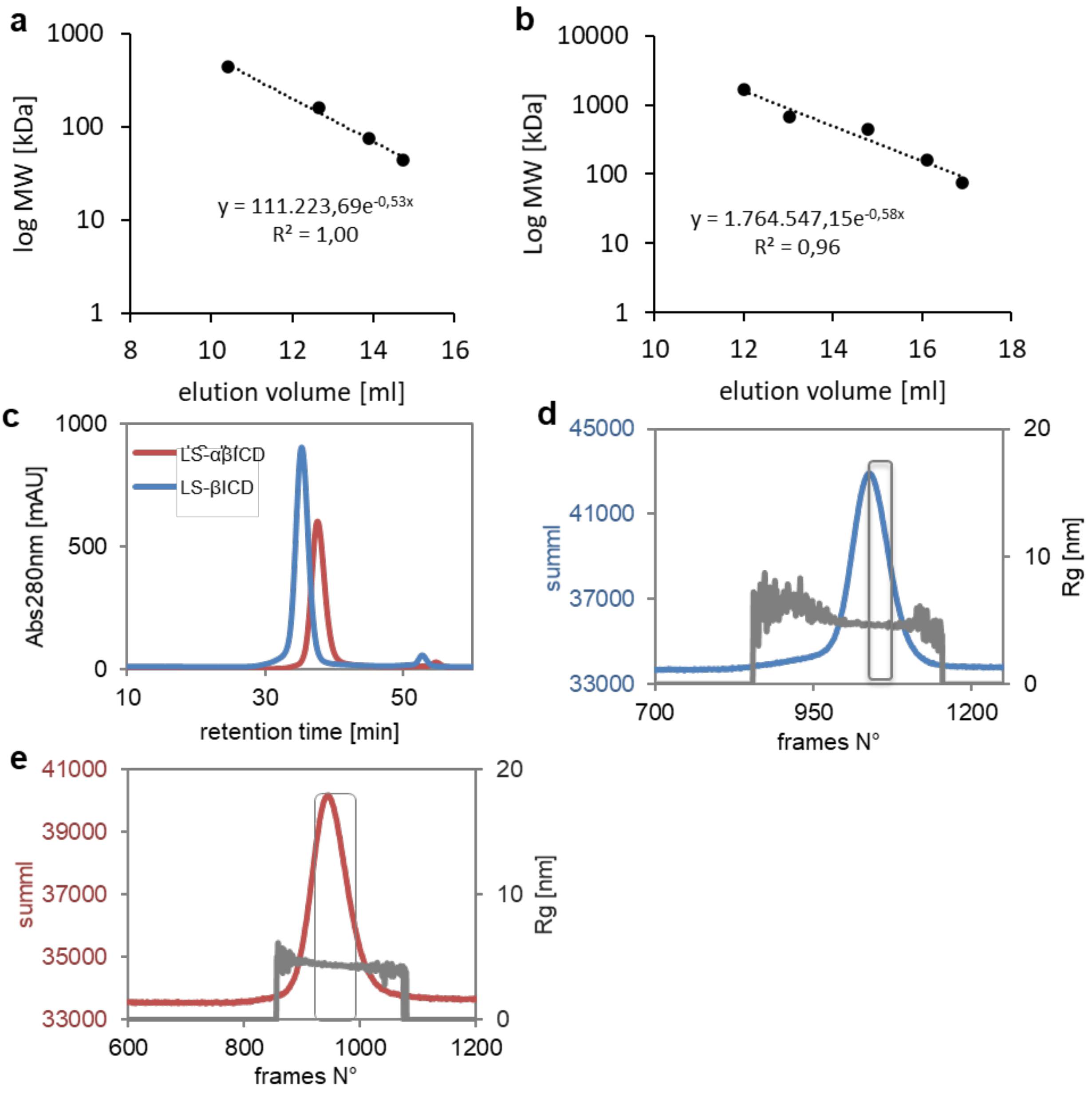


**Figure S7. Standard calibration curves for analytical SEC and SEC-coupled collection of SAXS-data. (a)** Superdex 200 10/300 GL, **(b)** Superose 6 10/300 increase. Ovalbumin (44 kDa), Covalbumin (75 kDa), Aldose (158 kDa) Ferritin (440 kDa). Thyroglobulin (669 kDa) and respiratory chain supercomplex I/III2/IV (1.7 MDa) were added for Superose 6 10/300 calibration served as standard proteins. **(c)** UV-280nm absorbance profiles of LS-βICD (blue) and LS-αβICD (red) collected during corresponding SAXS measurements. **(d, e)** Rg-values and raw scattering intensities (SummI) measured during SEC-runs of LS-βICD (d) and LS-αβICD (e) , respectively. Grey boxes indicate frames, which were used for the data processing.

**Supplemental Tables**

**Table S1. Diameters of pGLICs compared to that of LS.**

| Pentamer | Averaged diameter [Å] [6] |
|---|---|
| nAchR[1] | 72 |
| $GABA_AR$ β3 [2] | 72 |
| GlyR $\alpha_1$ [3] | 71 |
| GlyR $\alpha_3$ [4] | 71 |
| LS [5] | 75 |

[1] (Unwin, 2005); [2] (Miller and Aricescu, 2014); [3] (Du et al., 2015); [4] (Huang et al., 2015); [5] (Meining et al., 2000); [6] Diameters of pentameric assemblies were determined by PYMOL using the available crystal structures and were averaged from three measurements between different positions of the pentamer.

**Table S2. SAXS data collection parameters**

| | |
|---|---|
| Detector | PILATUS 1 M |
| Detector distance (m) | 2.867 |
| Beam size (µm x µm) | 700 x 700 |
| Wavelength (Å) | 0.99 |
| Sample environment | Quartz glass capillary, 1 mm ø |
| s range ($nm^{-1}$)‡ | 0.025–5.0 |
| Temperature (°C) | 4 |
| Exposure time per frame | 10 x 1s in static mode, 1.5 s continuously in online mode |
